# Distance-guided protein folding based on generalized descent direction

**DOI:** 10.1101/2021.05.16.444345

**Authors:** Liujing Wang, Jun Liu, Yuhao Xia, Jiakang Xu, Xiaogen Zhou, Guijun Zhang

## Abstract

Advances in the prediction of the inter-residue distance for a protein sequence have increased the accuracy to predict the correct folds of proteins with distance information. Here, we propose a distance-guided protein folding algorithm based on generalized descent direction, named GDDfold, which achieves effective structural perturbation and potential minimization in two stages. In the global stage, random-based direction is designed using evolutionary knowledge, which guides conformation population to cross potential barriers and explore conformational space rapidly in a large range. In the local stage, locally rugged potential landscape can be explored with the aid of conjugate-based direction integrated into a specific search strategy, which can improve exploitation ability. GDDfold is tested on 347 proteins of a benchmark set, 24 FM targets of CASP13 and 20 FM targets of CASP14. Results show that GDDfold correctly folds (TM-score ≥ 0.5) 316 out of 347 proteins, where 65 proteins have TM-scores that are greater than 0.8, and significantly outperforms Rosetta-dist (distance-assisted fragment assembly method) and L-BFGSfold (distance geometry optimization method). On CASP FM targets, GDDfold is comparable with five state-of-the-art methods, namely, Quark, RaptorX, Rosetta, MULTICOM and trRosetta in the CASP 13 and 14 server groups.

## 1 Introduction

Protein structure prediction has witnessed remarkable progress recently due to advances in protein folding from scratch driven by accurate prediction of the inter-residue distance [1–5]. Distance encodes the three-dimensional structure of a protein through paired residue spatial information, which can be converted to physical constraints to drive the folding of the protein from scratch [6–9]. Recent works on distance prediction and distance-based protein structure modeling have attracted wide attention [10–12], particularly in some top-performing groups in latest CASPs.

Fragment assembly [13,14] based conformational sampling approach is one of the primary paradigms for tackling folding problems. Constructing knowledge-based and physics-based energy potentials, low-energy basins are sampled by fragment assembly where distance is usually used as an energy term or constraint to assist conformational sampling. Rosetta [15] and QUARK [16] are two advanced de novo protein structure prediction methods based on fragment assembly. In CASP13, a composite energy function that adds contact restraints is designed by QUARK, which is used to guide the fragment assembly process through replica-exchange Monte Carlo simulation [17]. In the recent CASP14, predicted distance is integrated into the complex energy function in QUARK to guide protein folding [18]. Fragment assembly converts dihedral continuous space into discrete experimental fragment space, which can effectively capture the local propensities of known protein structures and reduce search space. However, promising conformations in the continuous search space may not be sampled sufficiently because of the limited fragment library [19–22]. In addition, the conformational space vastly increases with increasing protein length, and such increase leads to unsatisfactory prediction accuracy and expensive calculation cost for large proteins.

Recently, distance-restrained geometry optimization methods have led to a paradigm shift in protein structure prediction [23,24]. RaptorX [25], the first to predict discrete distances, uses the mean and variance of a predicted distance distribution to calculate lower and upper bounds for atom-atom distances and then feed calculated values to Crystallography and NMR System (CNS) [26] to fold proteins. MULTICOM [27] uses simulated annealing optimization implemented in CNS to build tertiary structure models by satisfying torsion angle, atom-atom distance, and hydrogen-bond restraints as good as possible. AlphaFold [24,28] converts full distance distribution as a smooth statistical potential function, and then the gradient descent L-BFGS [29] is used in optimizing directly the three-dimensional structures of a protein. In trRosetta [30], predicted distance and orientation are used as constraints, and a smooth potential is constructed through spline interpolation. The energy-minimization protocol of Rosetta is used to generate structure models guided by the potentials of predicted inter-residue orientations and distances.

The prevalent geometry optimization method optimizes a defined distance potential directly with gradient descent. The common problem of multiple minima persists, even for a reasonably accurate distance potential. Gradient-based optimization is effective in finding the nearest local minimum in a potential landscape, but it will not generally locate the global minimum [20]. Some attempts have been made to restart folding trajectories several times, given the sensitivity to initial conformations for gradient-based methods [28,30]. Combined with the inherent difficulty of gradient derivation, it may result in the low computational efficiency of distance geometry optimization method. Therefore, inappropriate search strategies may cause stagnation because of over exploration or premature convergence due to over exploitation. Designing two stages with the cooperation of distinct generalized descent directions may be effective. The former stage aims to locate promising regions rapidly, whereas the later stage encourages local exploitation for enhancing convergence.

In this article, we propose a distance-guided protein folding algorithm named GDDfold, which is based on generalized descent direction. Under an evolutionary algorithm framework [31,32], GDDfold integrates generalized descent directions into distinct search strategies corresponding to two stages. In the global stage, the random-based descent direction is designed using evolutionary knowledge, which guides conformations cross potential barriers and explore promising regions rapidly. As to the local stage, a locally rugged potential landscape can be explored with the aid of a conjugate-based direction designed based on the local convexity theory [33,34]. A search strategy with the cooperation of different directions is developed in the local stage to prevent premature convergence and improve exploitation ability. Two stages are switched by an evolutionary state estimation mechanism. Experimental results show that GDDfold can effectively prevent trapping into local minima and significantly improves the prediction accuracy of the distance geometry optimization method.

## 2 Materials and methods

### 2.1 Overview

The pipeline is shown in Figure 1. The sequence and the inter-residue distance are used as inputs, and GDDfold finally outputs the predicted three-dimensional structure of a target sequence. GDDfold is developed on the framework of evolutionary algorithm. First, the initial population is generated through random dihedral angle perturbation. In the global stage, random-based descent directions for each conformation are constructed using evolutionary knowledge of population interaction, which enables a population to cross potential barriers and quickly locate promising regions. In the local stage, conjugate-based descent directions are constructed using population historical information to quickly converge to the local optimal in the promising region which was explored before. The two stages are switched by evolutionary state estimation mechanism. Finally, the lowest-potential conformation is selected from the final population as the prediction model.

**Figure 1.**
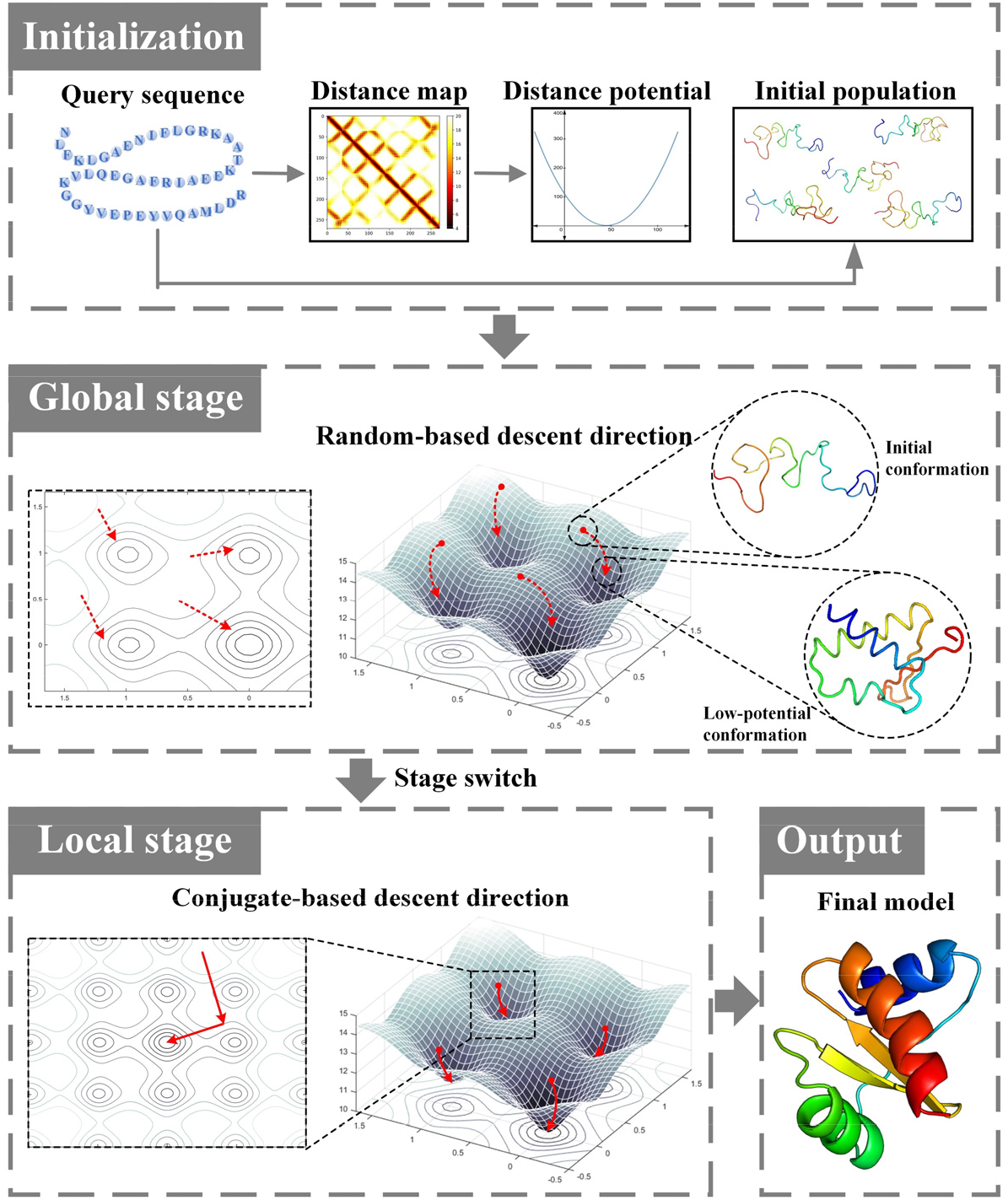
Pipeline of GDDfold. (1) Initialization. The initial conformations and the distance-based potential function are constructed. (2) Global stage. Low-potential conformations are generated by searching along random-based descent directions (dashed red arrows) from initial conformations. The search steps on the contour (left part) and surface (right part) of potential are illustrated. (3) Local stage. The low-potential conformation is further enhanced along the conjugate-based descent direction (solid red arrows). The search steps are also shown (left part is contour and right part is surface). (4) Output. Final prediction model.

In general, inter-residue spatial distance information is converted to an energy potential for the ab initio folding process. Distance probabilities are estimated for discrete distance bins. Here, the potential is constructed with a Gaussian distribution function. The distance potential is created from the negative log likelihood of the distances and is defined as follows:

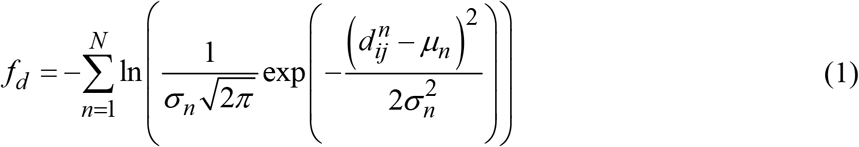

where *N* is the number of effective inter-residue distance; *i* and *j* are the residue indexes of the *n*-th distance; 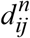 is the real distance between *C_β_* (*C_α_* for glycine) of residues *i* and *j* in the evaluated conformation; and *μ_n_* and *σ_n_* are the mean and standard deviation obtained by Gaussian fitting of the *n*-th inter-distance distribution, respectively.

### 2.2 Global stage

In the global stage, the early stage of search process, the conformations in a population are scattered in the different areas of a potential landscape. For an energy potential with multiple minima, designing a strategy with high speed and low computational efficiency is necessary to the exploration and discovery of more promising regions. Through the interaction and information exchange between the conformations in the evolutionary process, random-based descent direction can be constructed as follows:

First, the trial conformation 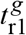 is obtained by searching along a random-based generalized descent direction starting from the selected basis conformation 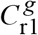.

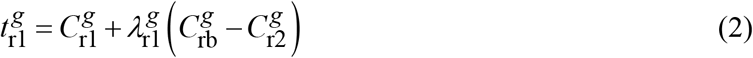

where 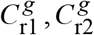, and 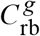 are the conformations chosen randomly from the population, and they differ from each other. 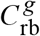 is a better conformation than 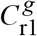 and 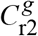, that is, the potential of 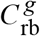 is lower. 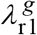 is the step size for 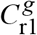, and the details are shown in the Supplementary materials.

The search procedure is illustrated in Figure 2. The core of the search procedure is to act the generalized descending direction composed of two randomly selected conformations on the randomly selected basis conformation. The interactive information is extracted from the positional relationship between the conformation in a population to guide the search process, enabling the population easy to search the conformational space in a large range and cross the potential barriers.

**Figure 2.**
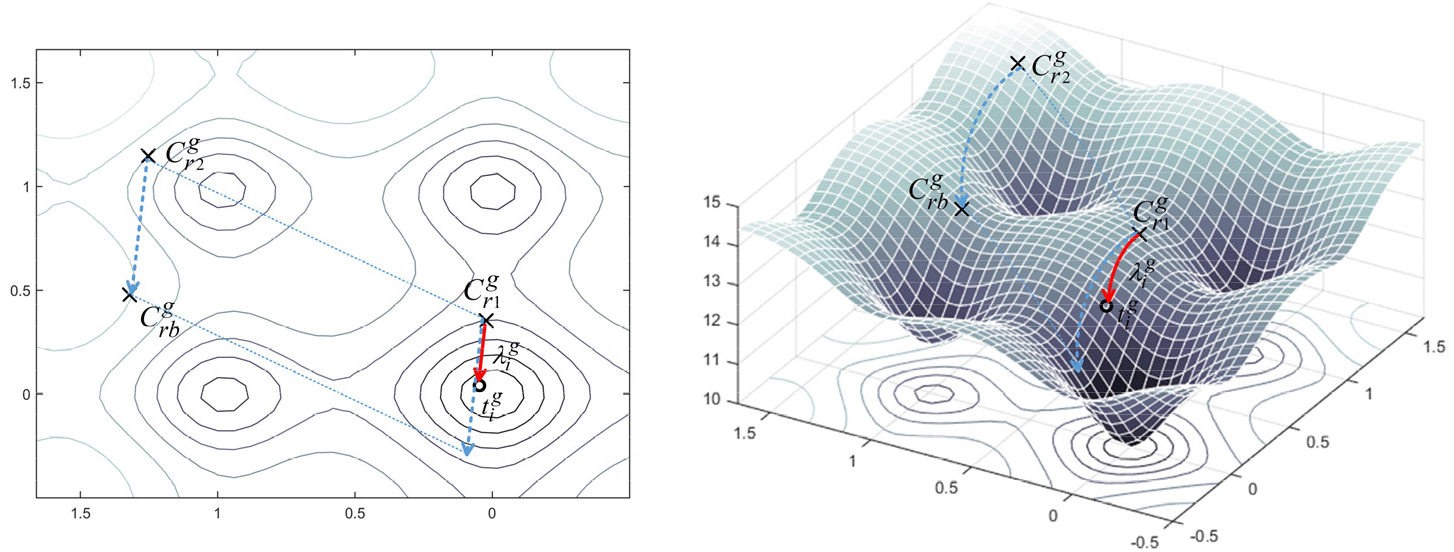
Schematic of search step with random-based descent direction. The crosses represent the conformations of generation *g,* and the circle represents the trial conformations 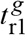. The random-based descent direction constructed by 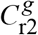 and 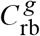 (marked in dash blue arrow) acts on the basis conformation 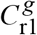. 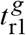 is obtained by searching along the above direction with step size 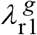. The actual search path is marked in solid red arrow.

For the detection and preservation of these promising regions, the crowding strategy is used to update population. After the replacement of the nearest parent conformation 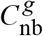 with the generated conformation 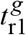, similar conformations tend to move closer to each other, whereas different conformations tend to move away from each other. The crowding selection operation can be expressed as follows:

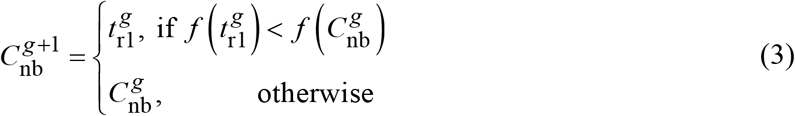

where 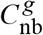 is determined by the computation of dihedral angle similarity. Given the conformation *C_m_* and *C_n_*, the similarity metric between them is defined as

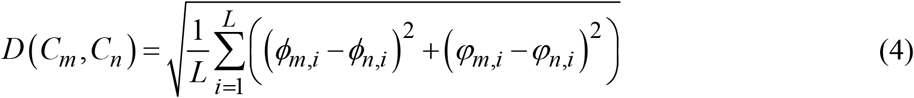

where *L* is the length of the protein sequence; *i* is the residue index; *φ_m,i_* and *φ_n,i_* are the dihedral angles about C-N-Cα-C for the *i*-th residue of *C_m_* and *C_n_*, respectively; and *φ_m,i_* and *φ_n,i_* are the dihedral angles about N-Cα-C-N for the *i*-th residue of *C_m_* and *C_n_*, respectively.

The above steps are performed *NP* times per generation, where *NP* is the population size. The basis conformations selected each time should be different from each other in order that each conformation in the population can be used as a starting point for searching along a random-based descent direction.

### 2.3 Local stage

As the search process proceeds, the population of conformations may converge to several locally or globally optimal regions. The local landscape surface that appears smooth is actually rugged, as shown in the Figure 3, and thus gradient descent methods easily fall into small local traps though the region containing the optimal solution have found. The reason is that the methods are highly dependent on the initial location because of the influence of the gradient direction. In this case, a search strategy adopted in the local stage should ensure that promising regions explored in the global stage are further exploited and convergence ability is enhanced.

**Figure 3.**
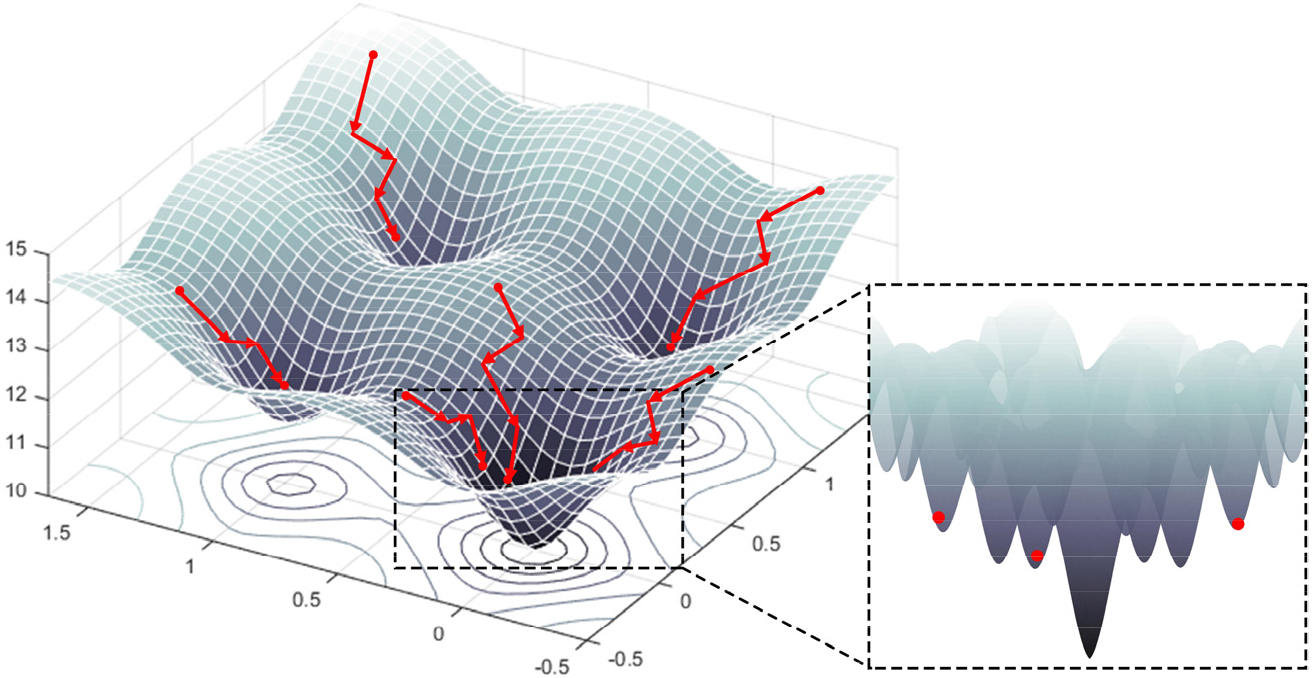
Schematic of one case in the local stage. For traditional gradient descent method, the conformations (red dots) search along the gradient directions (red arrows) on the surface of a potential, and the final results are highly dependent on initial conformations. When one of the regions containing minima (dashed box) are zoomed, the seemingly smooth surface is locally rugged, and thus traditional gradient descent method make it easy for conformations (red dots) to fall into small local traps.

Based on the local convexity theory [33,34], the objective function can be approximated by a quadratic function when near the minimum. Inspired by conjugate gradient method [35], a conjugate-based direction without gradient calculation is designed in the local stage. Two descent directions that are conjugate to each other are determined first. The search procedure is illustrated in Figure 4.

**Figure 4.**
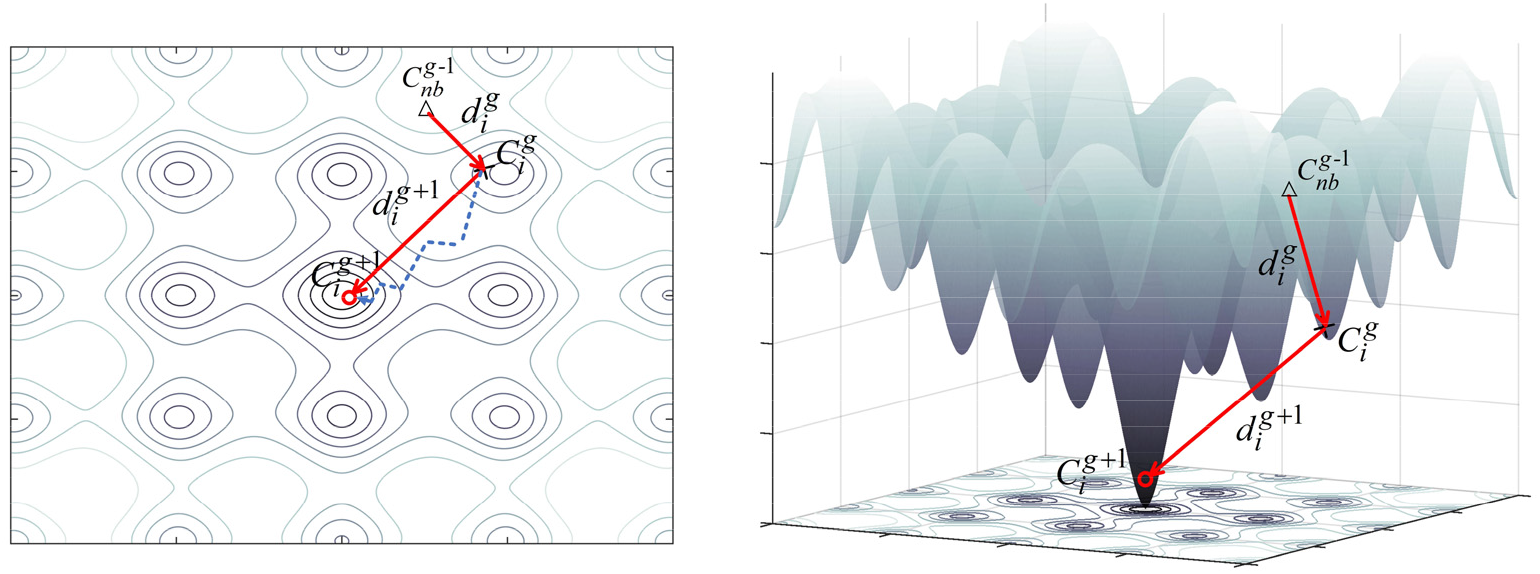
Schematic of search step with conjugate-based descent direction. 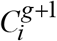 is obtained by searching from 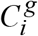. The triangles, crosses, and circles represent the conformations of generation *g*-1, *g* and *g*+1, respectively. For target 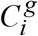, the solid red arrow denotes the local descent direction 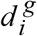 and the conjugate descent direction 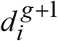. The dashed blue arrow represents the quasi-newton direction.

For target conformation 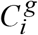, let 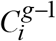 be the parent conformation of 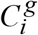 and 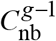 be the conformation closest to 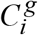 at the *g*-1th generation. The first direction 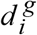 named local descent direction of 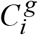 can be defined as follows:

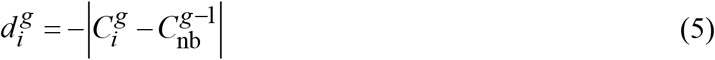

Therefore, the second direction 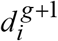 named conjugate descent directions of 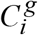 which is orthogonal to the local descent direction can be designed as follows:

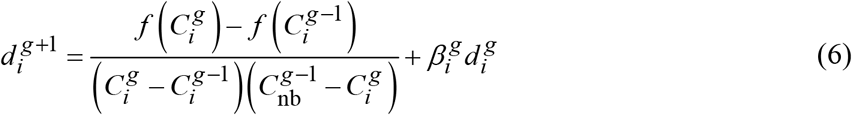

where 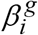 is the coefficient factor of combined search direction which can be calculated by

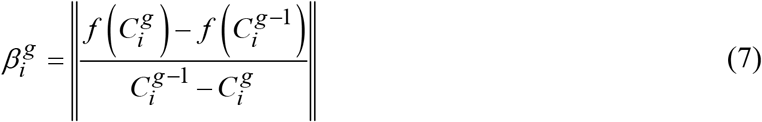

Finally, the offspring conformation 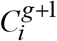 is obtained by searching along the conjugate descent direction from target conformation 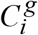 according to the following equation.

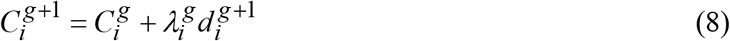

Considering the inherent greed of the conjugate method, the local random-based descent direction is used in the implementation of one-step search after searching from 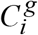 along the conjugate descent direction, which can realize the adjustment of conjugate direction of a small range to overcome the local barrier. Here, the target conformation is selected as the basis conformation, and one-to-one-spawning selection [36] is adopted. The cooperation of different directions in the above search strategy not only ensures the rapid convergence of the algorithm along the conjugate direction but also escapes small local traps through random direction.

### 2.4 Stage switch

Variations in search behaviours and population distributions during the search process allow search dynamics to induce basin-to-basin transfer, where trial conformations may traverse from one attraction basin to another one [37,38]. The scattered or converged feedback information of the population search behaviours is useful in constructing an entropy metric which reveals the trend of the movement of conformations in the search space to control the stage switching more objectively.

After crowding operation with several iterations in the global stage, *K* stable states can be obtained which correspond to different solution subspaces. In line with the temporal ordering of state transition of conformations at the two adjacent generations, a transition matrix 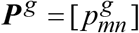 can be constructed, where 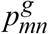 represents a transition probability between any given pair of states *m* and *n*.

Then, the entropy across all possible transitions can be calculated by summing the Shannon entropy for each conformation transition at each iteration.

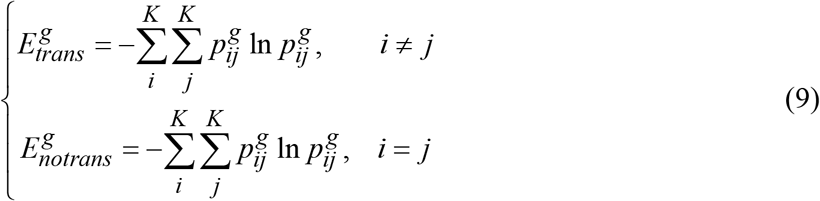

where 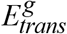 and 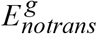 is denoted as the transition entropy and no transition entropy of the *g*th generation.

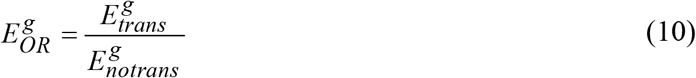

where 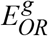 is odds ratio of 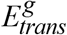 to 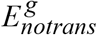.

After the evolutionary state is estimated, the stage switching more pertinent for conformations can be adaptively controlled to match the requirements of different stages. At the beginning of the search process, the value of 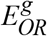 is greater than zero, and the algorithm is in the global stage. As search progresses, 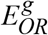 gradually decreases. Once the value of 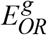 is equal to zero, the algorithm switches to the local stage. The details of stage switching mechanism are shown in the Supplementary materials material.

## 3. Result

### 3.1 Experiment settings and performance evaluation

The performance of GDDfold is tested on 347 non-redundant benchmark proteins systematically selected from the PDB and 44 FM targets from CASP13 and CASP14 proteins. The lengths of benchmark proteins range from 52 residues to 199 residues, with 30% sequence identity to each other. In SCOPe 2.07 [39], which have 243,819 proteins with known structures, a total of 11,198 non-redundant proteins are clustered by CD-HIT [40] with a 30% sequence identity cutoff. Afterward, 2,481 proteins are obtained after the exclusion of the multidomain proteins and limiting of protein length to 50-200. Eventually, 347 proteins are randomly selected from the 2,481 remaining proteins which vary in native fold topologies, including α, β, and α/β. For the 44 FM CASP proteins, 24 target proteins are from CASP13 and 20 target proteins are from CASP14. Detailed information of benchmark proteins and FM targets of CASPs is shown in Supplementary materials Table S2-S5, respectively.

GDDfold is a population-based structure prediction method, population size of *NP* = 100. We allow maximal generations of *G_global* = 100 for the global stage and *G_local* = 100 for the local stage. Two well-known structural quality measures are used in assessing the similarity of the predicted conformation and a reference conformation, generally the native structure. One is the root mean square deviation (RMSD) [15,41], which is defined as the average distance between Cα atomic coordinates after optimal rigid body superposition. The smaller RMSD means the smaller deviation and the better model accuracy. The other one is the template modeling score (TM-score) [42,43] which focuses on the measure of the global topology similarity and is robust to local structural variations. TM-score is length-independent of chain and ranges in (0,1], where higher value reflects better folding accuracy. Meanwhile, a TM-score ≥ 0.5 represents correctly folded models.

### 3.2 Comparison with distance geometry optimization method L-BFGSfold

To study the performance between GDDfold and one distance geometry optimization method, GDDfold is compared with L-BFGSfold, potential minimization implemented by L-BFGS, on the 347 proteins of a benchmark set. L-BFGSfold adopts the same evolutionary algorithm framework, which is integrated Rosetta’s quasi-newton-based energy minimization (Minmover) algorithm into. For fairness, GDDfold and L-BFGSfold use the same function formula of distance potential and the same initialization of conformation population.

The comparison of predicted results generated by GDDfold and L-BFGSfold is listed in Table 1 on all 347 benchmark proteins, and the detailed results of each protein are presented in Table S2 of Supplementary materials. The average RMSD of GDDfold (5.62Å) is reduced by 13.41% relative to that of L-BFGSfold (6.49Å), and the average TM-score by GDDfold (0.680) is 4.29% higher than that of L-BFGSfold (0.652). GDDfold obtain models with TM-scores of >0.5, >0.6, >0.7 and >0.8, which account for 91.07%, 78.10%, 54.76% and 18.73% of the total 347 proteins, respectively. L-BFGSfold obtain models with TM-scores of >0.5, >0.6, >0.7 and >0.8, which account for 86.17%, 72.91%, 44.38% and 11.53% of the total 347 proteins, respectively.

**Table 1.** Predicted results of GDDfold and L-BFGSfold.

| Method | RMSD | TM-score | #TM≥0.5 | #TM≥0.6 | #TM≥0.7 | #TM≥0.8 | P-value | Sig |
| --- | --- | --- | --- | --- | --- | --- | --- | --- |
| GDDfold | 5.62 | 0.680 | 316 | 271 | 190 | 65 | NA | NA |
| L-BFGSfold | 6.49 | 0.652 | 299 | 253 | 154 | 40 | 4.31E-57 | - |

#TM ≥ 0.5, ≥ 0.6, ≥ 0.7, ≥ 0.8 are the number of the predicted model with TM-score ≥ 0.5, ≥ 0.6, ≥ 0.7, ≥ 0.8.
P-value and Sig (significance) are the results of the Wilcoxon signed rank test.

Figure 5 intuitively reflects the comparison of GDDfold with L-BFGSfold on the benchmark data set. Compared with L-BFGSfold, GDDfold achieves a lower RMSD on 287 of 347 proteins, accounting for 82.71%, and a higher TM-score on 338 of 347 proteins, accounting for 97.41%. After the same inter-residue distance and population are used, the prediction accuracy of GDDfold significantly improves compared to that of L-BFGSfold. The Wicoxon signed-rank test results in the last two columns of Table 1 confirm prove the results.

**Figure 5.**
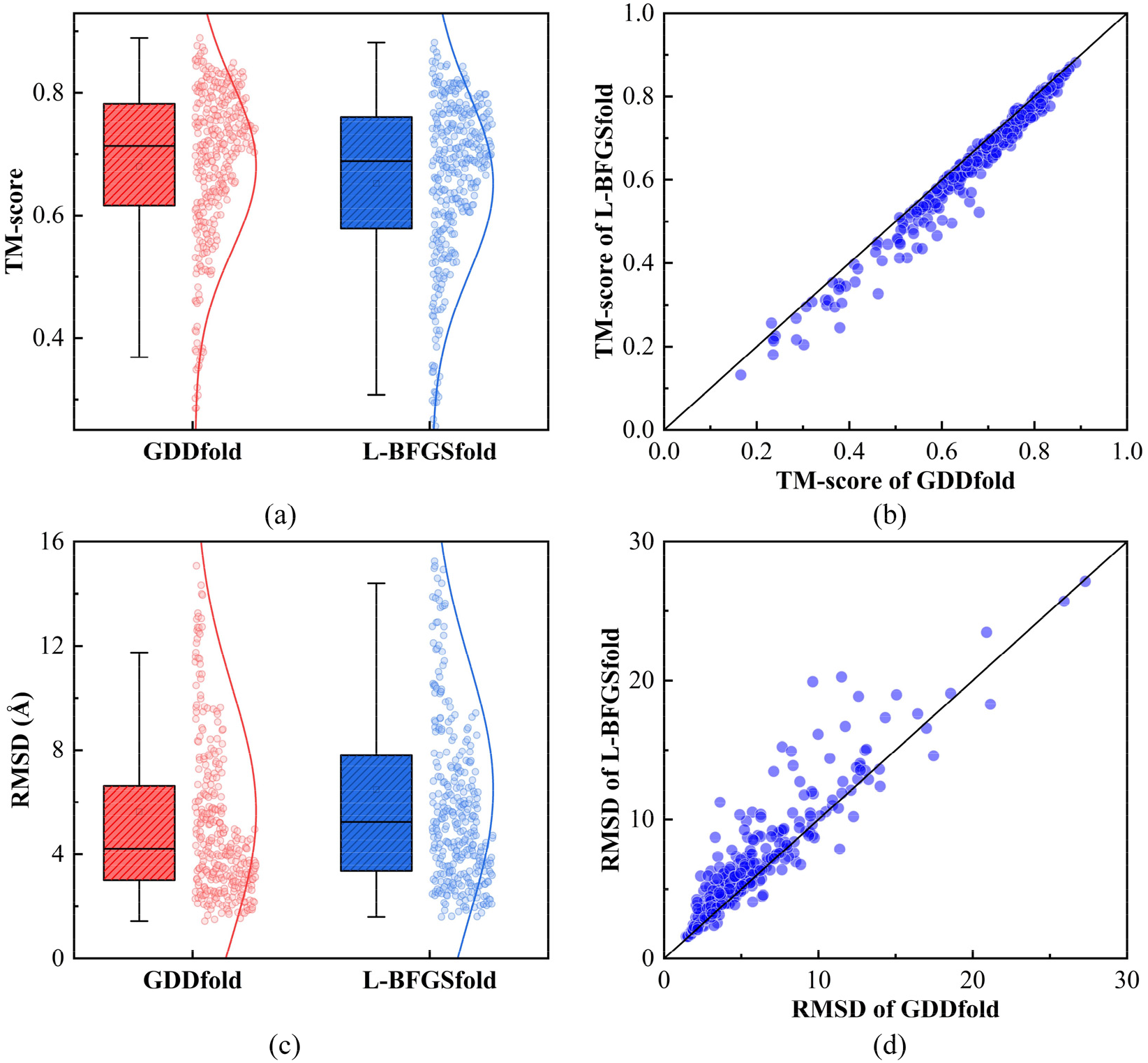
Comparison of the prediction performance of GDDfold and L-BFGSfold on all proteins. (a) Boxplot for TM-score of the predicted models by GDDfold and L-BFGSfold. (b) Head-to-head comparison between TM-score of the predicted models by GDDfold and L-BFGSfold. (c) Boxplot for RMSD of the predicted models by GDDfold and L-BFGSfold. (d) Head-to-head comparison between RMSD of the predicted models by GDDfold and L-BFGSfold.

### 3.3 Comparison with distance-assisted fragment assembly method Rosetta-dist

To investigate the performance between GDDfold and one distance-assisted fragment assembly method, GDDfold is compared with Rosetta-dist on the 347 proteins of the benchmark set. Rosetta-dist is implemented by Rosetta AbinitioRelax protocol, and the same distance potential as GDDfold is added to guide conformational sampling together with the energy function. A total of 500 candidate models are generated by running 500 independent trajectories, and then the candidate models are clustered by SPICKER [44]. The centroid structure of the first cluster is selected as the final model.

Comparison of predicted results generated by GDDfold and Rosetta-dist is listed in Table 2 on all 347 benchmark proteins, and the detailed results of each protein are presented in Table S2 of Supplementary materials. The average RMSD of GDDfold (5.62Å) is reduced by 17.84% compared to Rosetta-dist (6.84Å), and the average TM-score by GDDfold (0.680) is 25.69% higher than that of Rosetta-dist (0.541). GDDfold obtain models with TM-scores of >0.5, >0.6, >0.7, >0.8, which account for 91.07%, 78.10%, 54.76% and 18.73% of the total 347 proteins, respectively. Rosetta-dist obtain models with TM-scores of >0.5, >0.6, >0.7, >0.8, which account for 63.69%, 32.85%, 12.39% and 2.31% of the total 347 proteins, respectively.

**Table 2.**
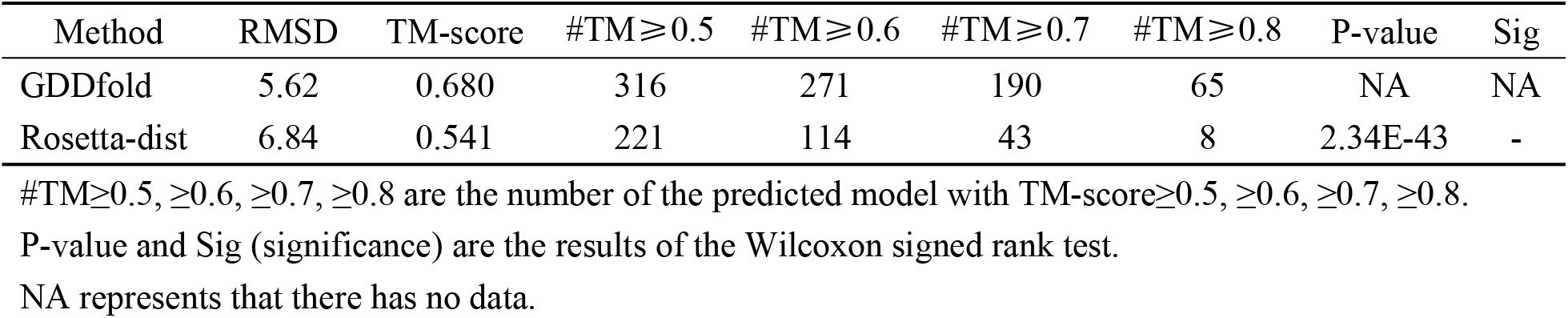
Predicted results of GDDfold and Rosetta-dist.

Figure 6 intuitively reflects the comparison of GDDfold with Rosetta-dist on the benchmark data set. Compared with Rosetta-dist, GDDfold achieves a lower RMSD on 253 of 347 proteins, accounting for 72.91%, and a higher TM-score on 291 of 347 proteins, accounting for 83.86%. GDDfold shows significant improvement in its prediction accuracy compared with Rosetta-dist (with P-values of <0.05).

**Figure 6.**
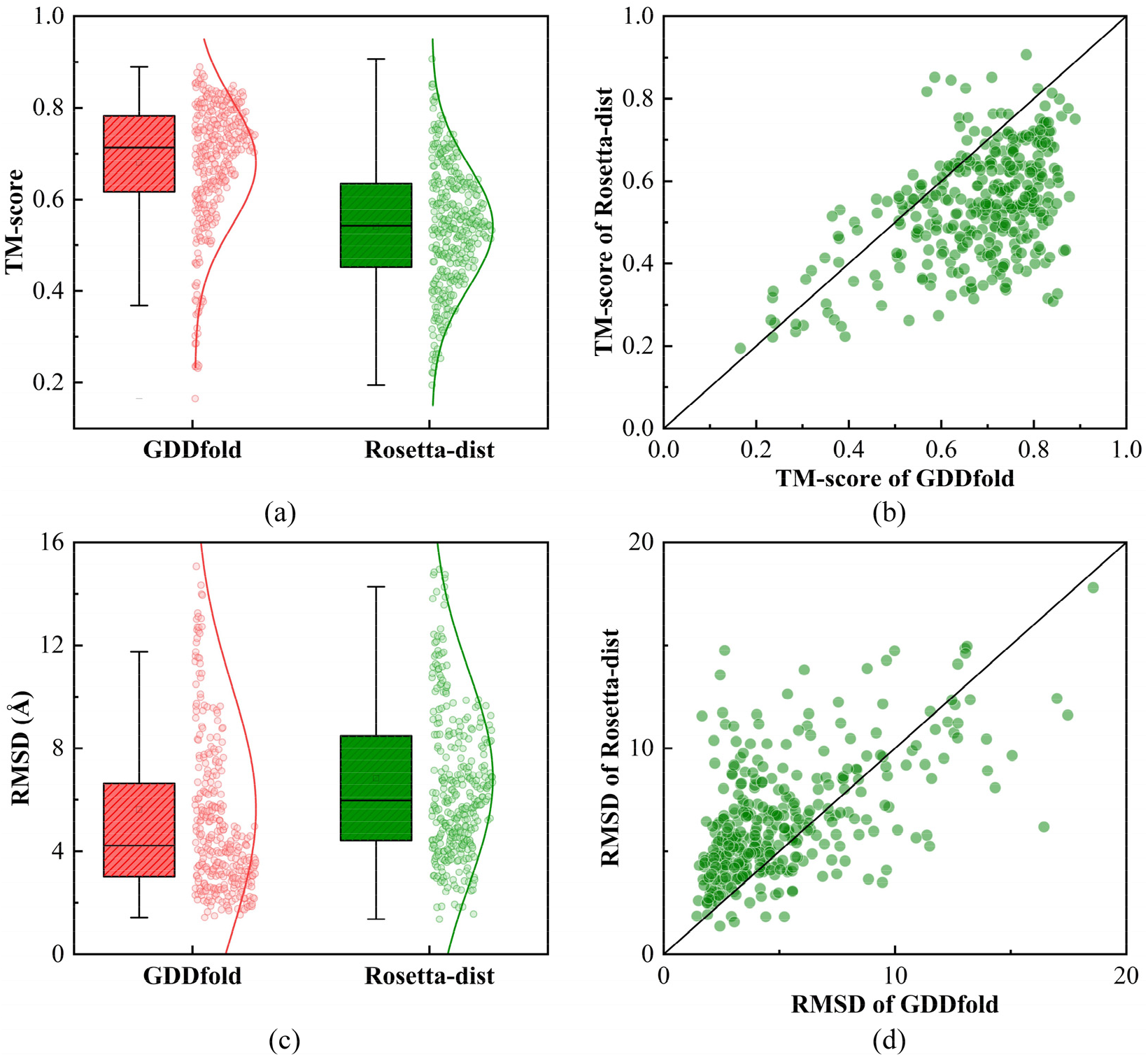
Comparison of the prediction performance between GDDfold and Rosetta-dist on all proteins. (a) Boxplot for the TM-scores of the predicted models by GDDfold and Rosetta-dist. (b) Head-to-head comparison between TM-score of the predicted models by GDDfold and Rosetta-dist. (c) Boxplot for RMSD of the predicted models by GDDfold and Rosetta-dist. (d) Head-to-head comparison between RMSD of the predicted models by GDDfold and Rosetta-dist.

### 3.4 Component analysis

For the identification of the impact of the two stages of GDDfold, component experiment is conducted on 347 proteins of the benchmark set. Two different variants, namely GDDfold_global and GDDfold_local, split from GDDfold as the two comparison algorithms. The difference is that only global stage is performed in GDDfold_global, whereas only local stage is implemented in GDDfold_local. The predicted results of GDDfold, GDDfold_global and GDDfold_local on the benchmark set are summarized in Table 3, and the detailed results of each protein are presented in Table S3 of Supplementary materials. The visual comparison of the performance of GDDfold, GDDfold_global, and GDDfold_local is illustrated in Figure 7.

**Table 3.**
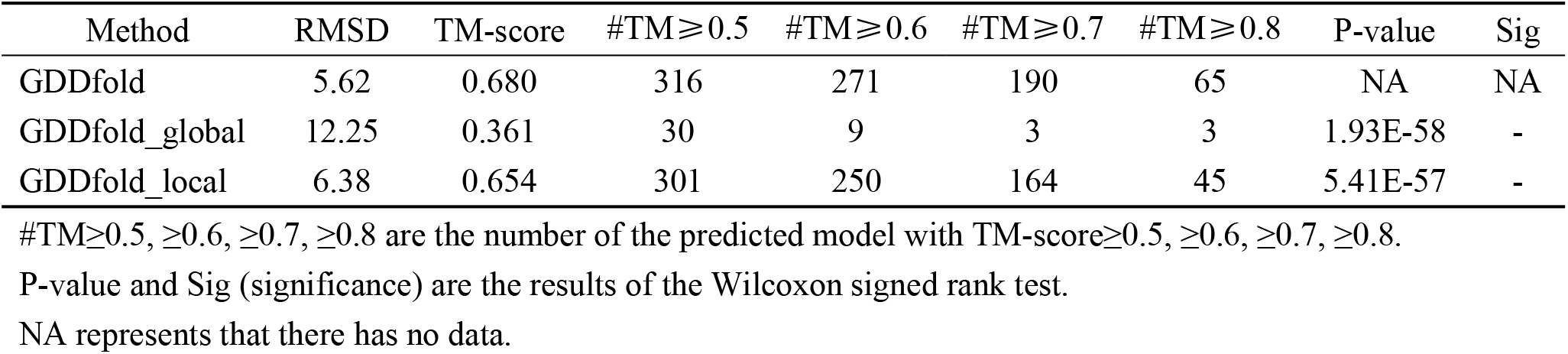
Predicted results of GDDfold, GDDfold_global and GDDfold_local.

| Method | RMSD | TM-score | #TM≥0.5 | #TM≥0.6 | #TM≥0.7 | #TM≥0.8 | P-value | Sig |
| --- | --- | --- | --- | --- | --- | --- | --- | --- |
| GDDfold | 5.62 | 0.680 | 316 | 271 | 190 | 65 | NA | NA |
| GDDfold_global | 12.25 | 0.361 | 30 | 9 | 3 | 3 | 1.93E-58 | - |
| GDDfold_local | 6.38 | 0.654 | 301 | 250 | 164 | 45 | 5.41E-57 | - |

#TM ≥ 0.5, ≥ 0.6, ≥ 0.7, ≥ 0.8 are the number of the predicted model with TM-score ≥ 0.5, ≥ 0.6, ≥ 0.7, ≥ 0.8.
P-value and Sig (significance) are the results of the Wilcoxon signed rank test.

**Figure 7.**
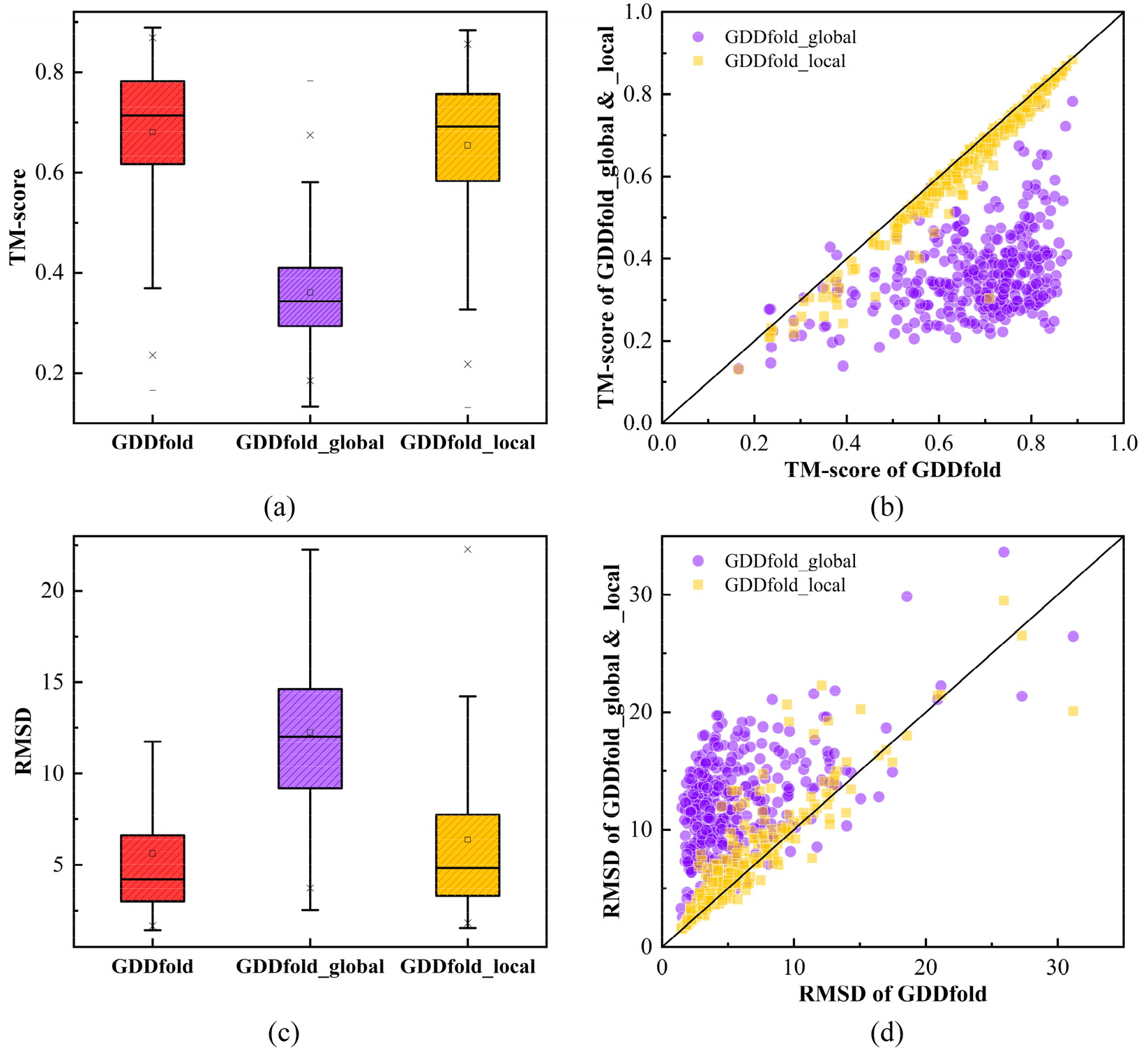
Comparison of the prediction performance between GDDfold, GDDfold_global, and GDDfold_local on all proteins. (a) Boxplot for TM-score of the predicted models by GDDfold, GDDfold_global, and GDDfold_local. (b) Head-to-head comparison between TM-score of the predicted models by GDDfold, GDDfold _global, and GDDfold_local. (c) Boxplot for RMSD of the predicted models by GDDfold, GDDfold_global and GDDfold_local. (d) Head-to-head comparison between RMSD of the predicted models by GDDfold, GDDfold_global and GDDfold_local.

Compared with GDDfold_global, the average RMSD of GDDfold decreases by 54.12%, and the average TM-score of GDDfold increases by 88.37%. The number of models correctly folded (TMscore ≥ 0.5) by GDDfold is 91.07%. The comparison results show that only the global stage of GDDfold does not work well. The random-based direction in global stage can guide the population to jump in a large range of conformational space. However, this advantage weakens in the later stage as the population converges, resulting in poorer search ability or even stagnation. Compared with GDDfold_local, the average RMSD of GDDfold is reduced by 11.91%, and the average TM-score of GDDfold is improved by 3.98%. The proportion of correctly folded models in GDDFold is 91.07%, which is more than that in GDDfold_local (86.74%). The comparison results show that only the local stage of GDDfold is insufficient. In the local stage, the population converges rapidly along the conjugate direction, which is fine-tuned by the cooperation of the local random-based direction. Compared with GDDfold_global, the conjugate-based direction of GDDfold_local plays an important role. However, compared with GDDfold, GDDfold_local may not find more promising regions because of the lack of a large-scale detection of the conformational space in the early stage. All the results in Table 3 show that the prediction accuracy of GDDfold is significantly better than each of its comparative versions. The cooperation of these two stages and the adaptive switching mechanism are effective.

### 3.5 Case study

Figure 8 shows the RMSD-iterations curve of L-BFGSfold and GDDfold on two proteins, and the conformation searched during certain iterations is marked on the figure. GDDFold is divided into two stages, and the impact of these stages on GDDfold is adequately reflected. In Figure 8 (a), as the number of iterations increases, the RMSD curve of L-BFGSfold dropps rapidly in the first few iterations, and then tends to be flat and nearly unchanged. The rapid convergence of L-BFGSfold in the previous iterations may be due to the effect of the quasi-newton direction, and the search stalls because of falling into the local optimum in the later period. In GDDfold, the first 100 iterations belong to the global stage and the latter 100 iterations belong to the local stage. A clear downward trend from the global stage switches to the local stage is indicated. In the global stage, the RMSD curve slowly declines and fluctuates possibly because the conformation population is scattered to explore the conformational space in a large range along random directions. When the stage is switched, a rapid decline in RMSD curve may be caused by switches from the random descending direction to the conjugate descending direction which has faster search speed. After entering the local stage, owing to the multiple promising basins explored in the global stage, the population continues to explore along the conjugate descending direction in each basin. Near the end of the algorithm and approaching the local minima, the step size at each iteration of the conformations gradually decreases, resulting in a flat RMSD curve. Compared with L-BFGSfold, GDDfold may explore more promising regions in the early stage, to a certain extent and prevent from falling into local traps, and finally find a conformation with a smaller RMSD. In addition, as shown in Figure 8 (b), the stage switching mechanism adaptively reduces the number of iterations of the global stage according to the current search situation, and thus the search efficiency of GDDfold is effectively improved.

**Figure 8.**
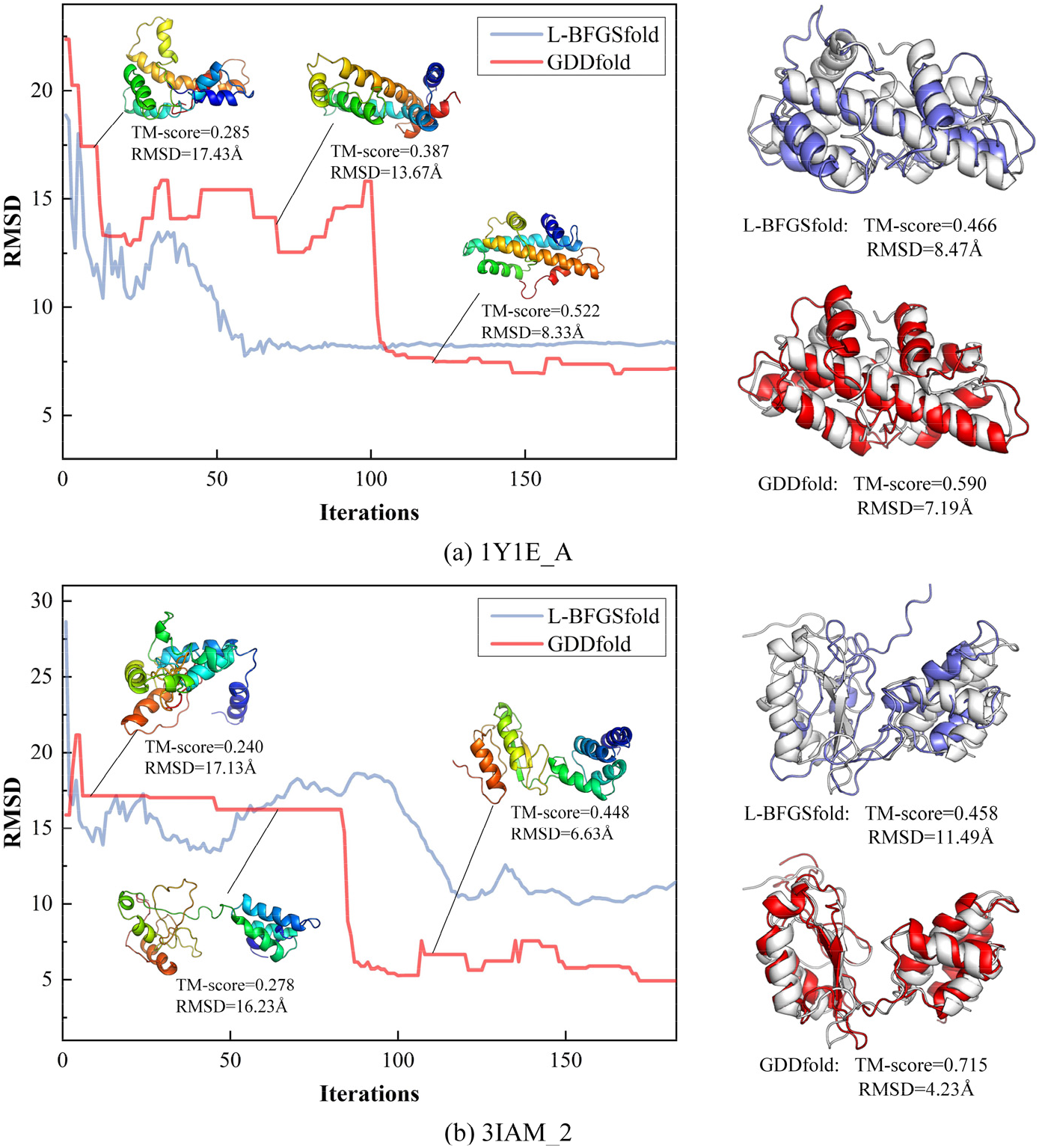
RMSD-iterations curves of L-BFGSfold and GDDfold on two proteins. The blue curve is L-BFGSfold and the red one is GDDfold. The predicted models compared with the native structure of lbfgs-fold and GDDfold are also marked. The structure of native, lbfgs-fold and GDDfold are marked in white, blue, and red, respectively.

### 3.6 Result of CASP13 and CASP14 Targets

GDDfold is compared with five methods of the server group in CASP 13 and 14, that is, QUARK, RaptorX, Rosetta, MULTICOM, trRosetta, which are state-of-the-art methods in the FM category of CASP experiments. The test proteins, 24 targets of CASP13 and 20 targets of CASP14, are downloaded from the CASP official website (https://predictioncenter.org/download_area/), as well as the results (best predicted model) of the above five methods. The names of the above methods are slightly different in CASP13 and 14. For example, trRosetta is named Yang-Server in CASP14. The details can be found in the Supplementary materials. In addition, GDDfold_relax is implemented where GDDfold plus FastRelax with distance constraints because only coarse-grained model is used in GDDfold compared with other methods. The results of GDDfold and other methods are shown in the Table 4 and Figure 9. Figure 10 shows the superimpositions of the target model predicted by GDDfold and the experimental structure. The detailed results are shown in Table S4, S5 of Supplementary materials.

**Table 4.** Prediction results of GDDfold, GDDfold_relax (GDDfold plus FastRelax with distance constraints), QUARK (QUARK for CASP13 and 14), RaptorX (RaptorX-DeepModeller for CASP13 and RaptorX for CASP14), Rosetta (BAKER-ROSETTASERVER for CASP13 and 14), MULTICOM (MULTICOM-CLUSTER for CASP13 and 14), and trRosetta (only Yang-Server for CASP14) on 24 FM targets from CASP13 and 20 FM targets of CASP14.

| Method in CASP13 | TM-score | #TM≥0.5 | P-value | Sig |
| --- | --- | --- | --- | --- |
| GDDfold | 0.544 | 15 | NA | NA |
| GDDfold_relax | 0.583 | 18 | 0.0034 | + |
| QUARK | 0.542 | 15 | 0.4567 | ≈ |
| RaptorX-DeepModeller | 0.516 | 14 | 0.1902 | ≈ |
| BAKER-ROSETTASERVER | 0.471 | 9 | 0.0063 | - |
| MULTICOM_CLUSTER | 0.417 | 6 | 0.0032 | - |
| Method in CASP14 | TM-score | #TM≥0.5 | P-value | Sig |
| GDDfold | 0.437 | 6 | NA | NA |
| GDDfold_relax | 0.465 | 9 | 0.1020 | ≈ |
| QUARK | 0.555 | 11 | 0.0249 | + |
| RaptorX | 0.447 | 7 | 0.5866 | ≈ |
| BAKER-ROSETTASERVER | 0.420 | 6 | 0.5564 | ≈ |
| MULTICOM_CLUSTER | 0.478 | 9 | 0.3951 | ≈ |
| Yang-server | 0.514 | 11 | 0.1771 | ≈ |

**Figure 9.**
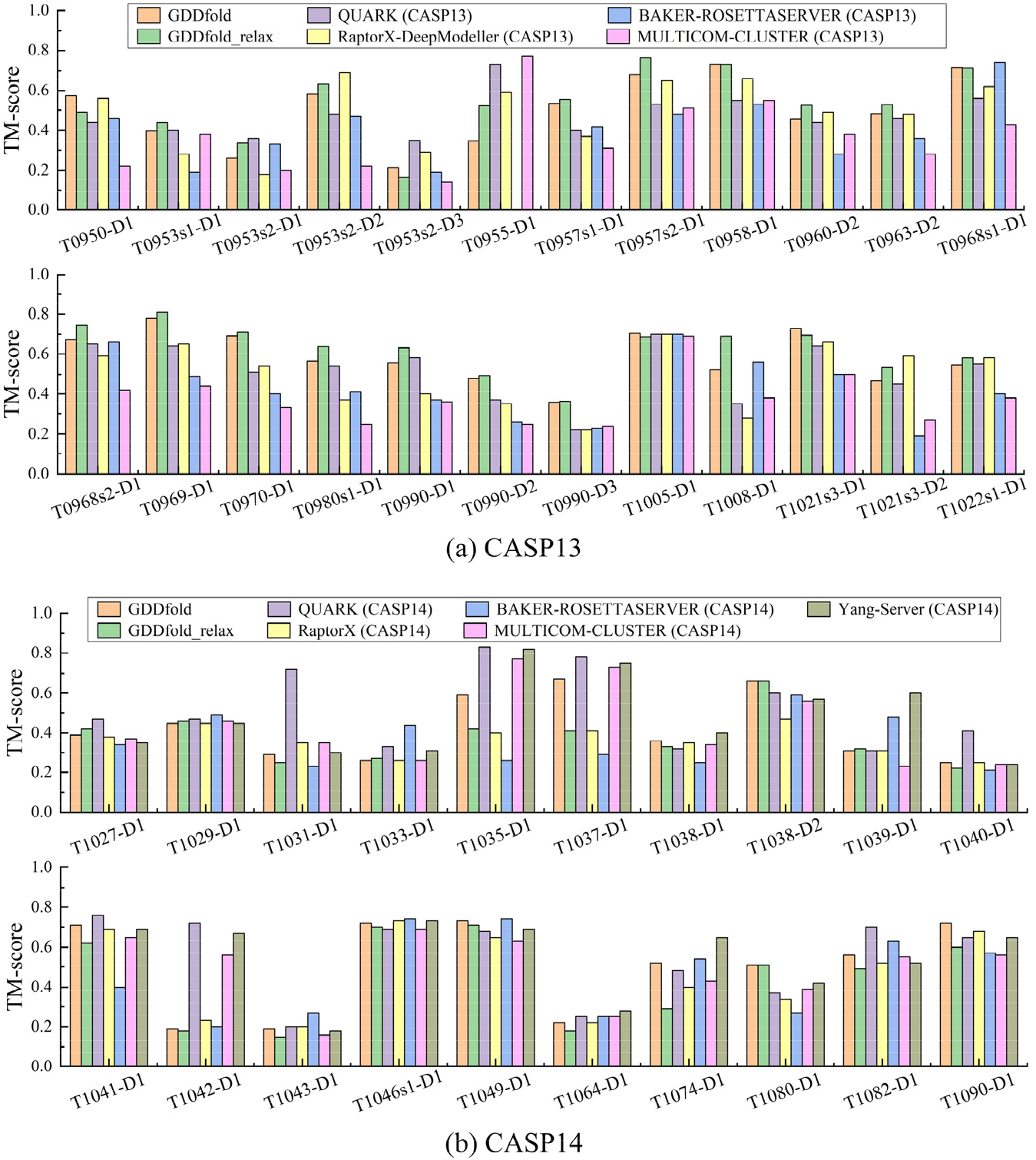
TM-score of the predicted model by GDDfold, GDDfold_relax (GDDfold plus FastRelax with distance constraints), QUARK (QUARK for CASP13 and 14), RaptorX (RaptorX-DeepModeller for CASP13 and RaptorX for CASP14), Rosetta (BAKER-ROSETTASERVER for CASP13 and 14), MULTICOM (MULTICOM-CLUSTER for CASP13 and 14), and trRosetta (only Yang-Server for CASP14) on 44 FM targets. (a) Results on the 24 FM targets of CASP13. (b) Results on the 20 FM targets of CASP14.

**Figure 10.**
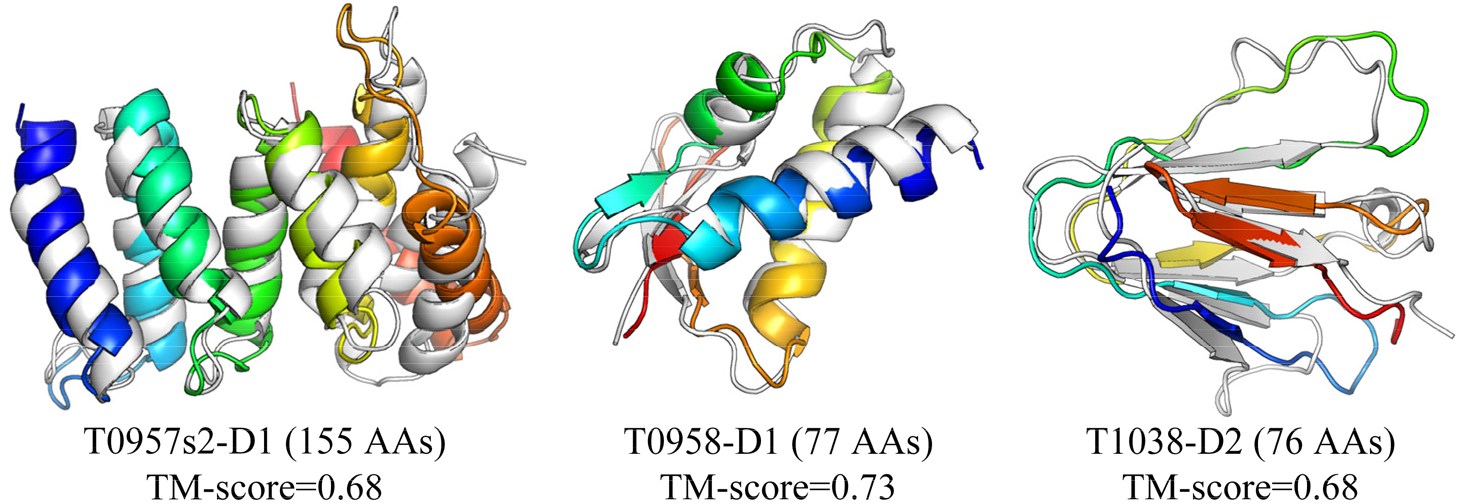
Superimposition between the predicted model (rainbow) by GDDfold and the experimental structure (white) for three FM targets (T0957s2-D1, T0958-D1, and T1038-D2) of CASP13 and CASP14.

On 24 CASP13 FM targets, the average TM-scores of the predicted model by GDDfold and GDDfold_relax are 0.544 and 0.583, respectively, which are better than the those of other methods. The results of QUARK (QUARK[17]), RaptorX-DeepModeller (RaptorX [25]), BAKER-ROSETTASERVER (Rosetta[45]), and MULTICOM_CLUSTER (MULTICOM[27]) are 0.542, 0.516, 0.471, and 0.417, respectively. GDDfold and GDDfold_relax obtain the highest TM-score on 3 and 10 targets compared with all methods, whereas QUARK, RaptorX-DeepModeller, BAKER-ROSETTASERVER, and MULTICOM_CLUSTER obtain the highest TM-score on 5, 4, 5, and 1 protein(s), respectively. As shown in Table 4, the non-parametric test of Wilcoxon with a significance level of 0.05 is used in single-problem analysis. The performance of GDDfold is equal to that of QUARK and RaptorX-DeepModeller, and better than that of BAKER-ROSETTASERVER and MULTICOM_CLUSTER. The FastRelax version GDDfold_relax further improves the performance of GDDfold.

On 20 CASP14 FM targets, the average TM-score of the predicted model by GDDfold is 0.437, which is better than that of BAKER-ROSETTASERVER (0.420). GDDfold_relax (0.465) improves GDDfold performance and further surpasses RaptorX (0.447). The results of QUARK, MULTICOM_CLUSTER and Yang-Server are 0.555, 0.478 and 0.513, respectively. The Wilcoxon test results show that GDDfold is inferior to QUARK, but no significant difference is observed between the performance of GDDfold and that of RaptorX-DeepModeller, BAKER-ROSETTA SERVER, MULTICOM_CLUSTER, and Yang-Server (with P-values of >0.05). In the CASP14 Abstract book, these latest servers in CASP14 use other information besides distance restraints. For example, QUARK utilizes information such as metagenome sequence databases (BFD, Mgnify, IMG/M), orientation, threading, and templates.

## 5. Conclusion

We propose a distance-guided protein folding algorithm based on generalized descent direction in this paper, named GDDfold. Under an evolutionary algorithm framework, two stages, namely the global and local stages, which are coupled with distinct generalized descent direction are performed. More promising regions are explored as rapidly as possible by random-based direction in the global stage. As to the local stage, the conjugate-based directions without gradient calculation are integrated into a specific search strategy, given that the computational complexity and convergence speed have an advantage over those of other optimization techniques. Two stages are switched by an evolutionary state estimation mechanism for the adaptive reduction of computational costs.

GDDfold is tested on 347 benchmark proteins and 44 CASP FM targets. Better than two representative techniques, namely fragment assembly and distance geometry optimization, GDDfold correctly folds (TM-score ≥ 0.5) 316 out of 347 proteins, and 65 models have TM-scores that are greater than 0.8. Component experiment and case study show the impact of two stages and the effectiveness of cooperation of generalized descent directions. On the latest CASP FM targets, GDDfold is also comparable with five state-of-the-art full-version methods in the CASP 13 and 14 server groups. The results show that GDDfold is a powerful protein folding method. Given that GDDfold is only a folding algorithm for coarse-grained models, relax and model quality assessment may be the next promising direction for improvement.

## Availability

The source code and executable are freely available at https://github.com/iobio-zjut/GDDfold.

