## Supplementary Material for "Distance-guided protein folding based on generalized descent direction"

### S1. Stage switch

The search dynamics in evolutionary algorithm induces basin-to-basin transfer, where trial solutions may traverse from one attraction basin to another one. These basins are defined as evolutionary states with respect to the partition of the solution space. In this way, the transition matrix based on evolutionary state is constructed using the historical evolutionary information across generations to describe the frequency of state transition. Subsequently, the entropy metric is proposed to estimate the extent that the population explores the solution space, which is mainly used for the dynamic division of the different stages.

1. Evolutionary state determination

The determination of evolutionary state consists of archiving, crowding, and clustering operations. The learn period is set to *LP*.

1) Archiving operation
   Search operations act on the target conformation and generate the trial conformation. The trial conformation is archived into a corresponding collection who representative conformation is the closest parent one to the trial conformation.
2) Crowding operation
   Through replacing the nearest parent conformation by generated trial conformation, similar conformations tend to move closer to each other, while different conformations tend to move away from each other.
3) Clustering operation
   Starting from the conformation with minimum potential value, the clustering step is performed with representative conformation of collection as the center and the farthest distance between conformations of collection as the radius in turn according to the ascending order. The process is completed when all the individuals are assigned to a corresponding cluster.

All conformations of the population perform the above operations within specified number of iterations. Denote these final clusters as the evolutionary states $S$ represented by center and $K$ as the number of evolutionary states. After iterations, the number $K$ tends to be stable.

2. Transition matrix

For $K$ evolutionary states, the count of states transition can be obtained using historical evolutionary information across generations to build the transition matrix $\boldsymbol{T}^g$. The element $t_{ij}^g$ in row $i$ and column $j$ in this matrix corresponds to the observed frequency of transitions from state $i$ at the $g$-1th generation to state $j$ at the $g$th generation. Therefore, each element $t_{ij}^g$ in the transition matrix $\boldsymbol{T}^g$ can be described as

$$t_{ij}^g = \frac{Z\left(S_i^{g-1}, S_i^{g-1}\right)}{N\left(S_i^{g-1}\right)} \tag{S1}$$

where $Z\left(S_i^{g-1}, S_i^{g-1}\right)$ is the number of state transition from state $i$ at the $g$-1th generation to state $j$ at the $g$th generation, and $N\left(S_i^{g-1}\right)$ is the number of conformations located in state $i$ at the $g$-1th generation.

Based on the transition matrix, the probability that the evolutionary process is undergoing a transition between any given pair of states can be estimated by

$$p_{ij}^{g} = \frac{t_{ij}^{g}}{\sum_{m}^{K}\sum_{n}^{K} t_{mn}^{g}} \tag{S2}$$

where $\sum_{i}^{K}\sum_{j}^{K} p_{ij}^{g} = 1$.

3. Switching mechanism based on entropy

The entropy across all possible transitions can be calculated by summing the Shannon entropy for each conformation transition at each iteration

$$\begin{cases} E_{trans}^{g} = -\sum_{i}^{K}\sum_{j}^{K} p_{ij}^{g} \ln p_{ij}^{g}, & i \neq j \\ E_{notrans}^{g} = -\sum_{i}^{K}\sum_{j}^{K} p_{ij}^{g} \ln p_{ij}^{g}, & i = j \end{cases} \tag{S3}$$

where $E_{trans}^{g}$ and $E_{notrans}^{g}$ is denoted as the transition entropy and no transition entropy of the $g$th generation.

$$E_{OR}^{g} = \frac{E_{trans}^{g}}{E_{notrans}^{g}} \tag{S4}$$

where $E_{OR}^{g}$ is odds ratio of $E_{trans}^{g}$ to $E_{notrans}^{g}$.

With the evolutionary state been estimated, we can adaptively control the stage switching more pertinent for conformations to match the requirements of different stage. At the beginning of the search process, the value of $E_{OR}^{g}$ is greater than zero, and the algorithm is in the global stage. As the search progresses, $E_{OR}^{g}$ gradually decreases. Once the value of $E_{OR}^{g}$ is equal to zero, the algorithm switches to the local stage.

4. Stage switching analysis

This section focuses on the effect of entropy-based evolutionary state estimation mechanism for GDDfold. In GDDfold, $E_{OR}^{g}$ is constructed based on the transition distribution of the current population to control the switching of global and local stages. Along with the population evolution, $E_{OR}^{g}$ of a target protein decreases rapidly in the early part of the global stage and the flow tends to be smooth in the latter part of the global stage, as shown in Figure S1. This phenomenon is due to the decreased population diversity, and the conformations with low energy exist in the next generation. In this case, the transition between different states gradually becomes reduced and the transition between their own states gradually becomes centralized, so that entropy odds metric declines correspondingly. Although the margins of $E_{OR}^{g}$ changes for target proteins are different from one another, all the curves almost level out in the end and the value is near to zero. In addition, the curves of $E_{OR}^{g}$ can indicate the adaptation of different protein search states and reduce the computational cost.

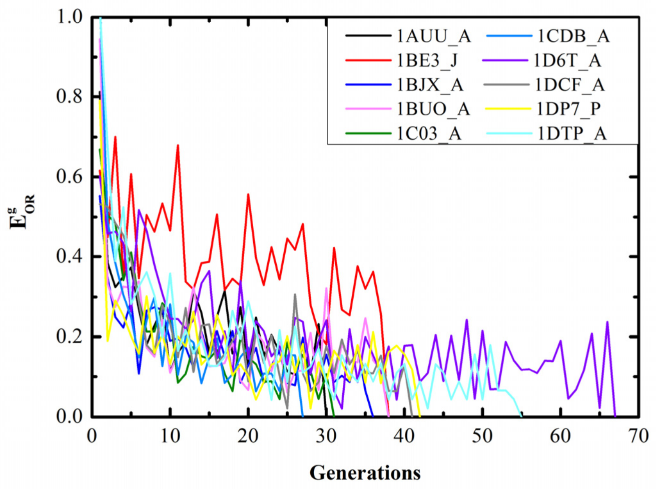


Figure S1. The entropy odds per generation for 10 proteins.

### S2. Overall implementation of GDDfold

The framework of GDDfold is described as Figure S1. The sequence and the inter-residue distance are used as input, and GDDfold finally outputs the predicted three-dimensional structure of the target sequence. GDDfold is developed on the framework of evolutionary algorithm. First, the initial population $P^g = \{C_i^g\}, i = 1,..., NP, g = 0$ is generated by random dihedral angle perturbation. After initialization, the $K$ evolutionary states are learned firstly through archiving, crowding, and clustering operations as shown in above section S2.

In the following iteration process, the conformations in population have state transition caused by the search behavior of the population. In line with the temporal ordering of state transition of conformations at the two adjacent generations, the state transition probability is calculated, and further the entropy based on transition matrix is calculated to observe the population dynamics. Based on the entropy metric, when to switch global stage into local stage can be estimated.

In global stage, the search operation with random-based direction and the crowding selection strategy are used to update population, which is followed by the update of entropy metric. Once the metric is zero, GDDfold automatically switches to the local stage. In local stage, the population converges rapidly along the conjugate direction which is fine-tuned by the cooperation of local random-based direction. Subsequently, the population is updated by one-to-one-spawning selection strategy. Finally, the lowest-potential conformation is selected from the final population as the prediction model.

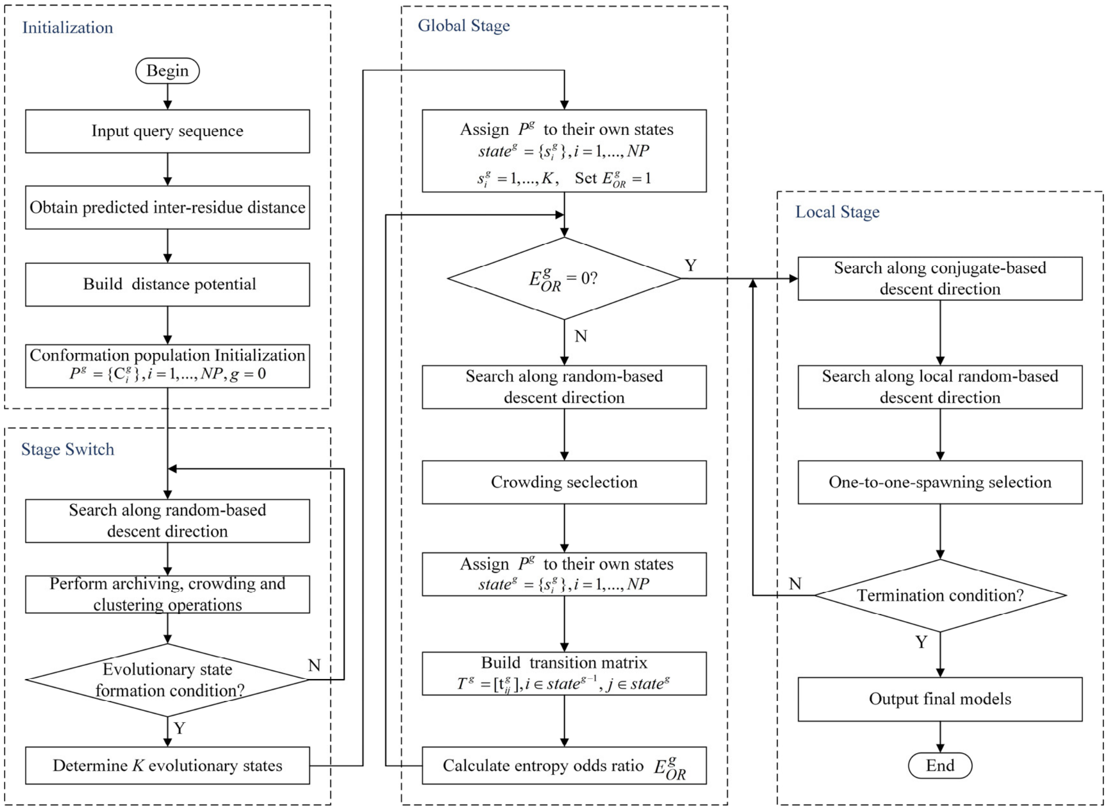


Figure S2. Framework of GDDfold.

### S3. Parameter setting of GDDfold

The parameters of MMpred are described in Table S1.

**Table S1.** The parameter descriptions in GDDfold.

| | | |
|---|---|---|
| Population size | *NP* | 100 |
| Generation of global stage | *G_global* | 100 |
| Generation of global stage | *G_local* | 100 |
| Initialization of step size | $\lambda$ | 0.5 |
| Learn period of stage switching | *LP* | 100 |

Parameter $\lambda_i^g$ needs to be described in detail. A simple self-adaptation strategy of $\lambda_i^g$ is designed for the requirements of different stages. The initial values for $\lambda_i^g$ is set to 0.5, and 0.5 is reassigned to $\lambda_i^g$ when the range of it exceeds.

In the global stage, the exploration state and the exploitation state are coexisted. This is because the conformations in the population are scattered at the early stage of evolution to explore extensively the search space. Due to the randomness, some individuals maybe find the local optimal region for exploitation. In this case, two different step sizes with distinct advantages are used to adapt the search feature in the global stage as follows:

$$\lambda_i^g = \begin{cases} \lambda_i^{g-1} + rand(0,0.1)E_{OR}^g, & \text{if } rand(0,1) \le E_{OR}^g \\ \lambda_i^{g-1} - rand(0,0.1)E_{OR}^g, & \text{otherwise} \end{cases} \tag{S5}$$

The large $E_{OR}^g$ caused by a case that the population is frequently transferred between several states, which indicates that the scope of conformational space is explored extensively. For this case, gradually larger step size with good exploration capability is more suitable. On the contrary, the population concentrates on some parts of the conformational space for exploitation when the value of $E_{OR}^g$ is small. Gradually smaller step size with good exploitation capability can be employed to detect the regions containing the minima.

In the local stage, a general line search technique, Armijo algorithm, is used to determine the step size $\lambda_i^g$.

**Table S2**. Prediction results of GDDfold, lbfgs-fold, and Rosetta-dist for 347 benchmark proteins.

| NO. | PDB | Length | Type | GDDfold | | lbfgs-fold | | Rosetta-dist | |
|---|---|---|---|---|---|---|---|---|---|
| | | | | RMSD | TMscore | RMSD | TMscore | RMSD | TMscore |
| 1 | 1A3A_C | 146 | α/β | 2.30 | 0.854 | 2.49 | 0.832 | 5.37 | 0.596 |
| 2 | 1A6L_A | 106 | α/β | 2.57 | 0.749 | 4.88 | 0.682 | 4.45 | 0.534 |
| 3 | 1A7D_A | 118 | α | 4.35 | 0.808 | 4.14 | 0.797 | 4.12 | 0.709 |
| 4 | 1A91_A | 79 | α | 4.95 | 0.461 | 5.01 | 0.451 | 4.43 | 0.522 |
| 5 | 1ABT_A | 74 | α/β | 4.79 | 0.576 | 6.25 | 0.552 | 4.26 | 0.495 |
| 6 | 1ABV_A | 105 | α | 3.04 | 0.698 | 3.04 | 0.689 | 3.04 | 0.720 |
| 7 | 1AHK_A | 129 | β | 5.79 | 0.531 | 5.97 | 0.522 | 5.59 | 0.539 |
| 8 | 1AK6_A | 174 | α/β | 6.03 | 0.670 | 6.81 | 0.662 | 6.29 | 0.534 |
| 9 | 1AKP_A | 114 | β | 3.07 | 0.712 | 4.67 | 0.659 | 9.13 | 0.402 |
| 10 | 1AP7_A | 168 | α | 3.37 | 0.761 | 5.25 | 0.741 | 4.23 | 0.691 |
| 11 | 1AUU_A | 55 | β | 2.12 | 0.680 | 4.26 | 0.638 | 3.06 | 0.590 |
| 12 | 1AX8_A | 130 | α/β | 4.06 | 0.687 | 4.39 | 0.641 | 9.42 | 0.416 |
| 13 | 1B4R_A | 80 | β | 2.33 | 0.743 | 2.43 | 0.723 | 3.62 | 0.573 |
| 14 | 1B4U_A | 132 | α | 11.38 | 0.597 | 7.88 | 0.562 | 5.78 | 0.637 |
| 15 | 1B8Q_A | 127 | α/β | 9.19 | 0.505 | 9.14 | 0.457 | 10.75 | 0.416 |
| 16 | 1BE3_J | 62 | α | 9.63 | 0.413 | 8.87 | 0.355 | 4.10 | 0.500 |
| 17 | 1BGF_A | 124 | α/β | 2.82 | 0.734 | 2.98 | 0.716 | 3.73 | 0.673 |
| 18 | 1BGY_J | 62 | α | 9.45 | 0.381 | 9.08 | 0.344 | 3.49 | 0.530 |
| 19 | 1BJX_A | 110 | α/β | 9.74 | 0.746 | 8.70 | 0.729 | 7.15 | 0.624 |
| 20 | 1BUO_A | 121 | α/β | 2.93 | 0.776 | 5.16 | 0.741 | 4.47 | 0.638 |
| 21 | 1C03_A | 163 | α/β | 11.58 | 0.609 | 12.74 | 0.601 | 8.53 | 0.556 |
| 22 | 1C41_A | 165 | α/β | 3.45 | 0.844 | 2.57 | 0.839 | 5.74 | 0.631 |
| 23 | 1C9F_A | 87 | α/β | 3.47 | 0.612 | 3.87 | 0.608 | 3.81 | 0.623 |
| 24 | 1CDB_A | 105 | β | 4.67 | 0.706 | 4.99 | 0.694 | 6.04 | 0.464 |
| 25 | 1CF7_B | 82 | α/β | 2.57 | 0.715 | 3.22 | 0.661 | 2.95 | 0.673 |
| 26 | 1CHC_A | 68 | α/β | 5.59 | 0.513 | 7.52 | 0.504 | 5.66 | 0.423 |
| 27 | 1CTO_A | 109 | β | 4.87 | 0.701 | 5.24 | 0.657 | 6.27 | 0.427 |
| 28 | 1CXZ_B | 86 | α | 5.22 | 0.621 | 5.84 | 0.585 | 1.83 | 0.845 |
| 29 | 1D6T_A | 117 | α/β | 4.60 | 0.753 | 4.90 | 0.748 | 5.12 | 0.587 |
| 30 | 1D8B_A | 81 | α | 2.46 | 0.715 | 2.67 | 0.691 | 3.39 | 0.591 |
| 31 | 1DBF_A | 127 | α/β | 4.02 | 0.802 | 5.61 | 0.767 | 5.58 | 0.526 |
| 32 | 1DCF_A | 133 | α/β | 3.21 | 0.813 | 4.64 | 0.799 | 3.97 | 0.751 |
| 33 | 1DL6_A | 58 | β | 11.74 | 0.378 | 16.70 | 0.351 | 10.91 | 0.403 |
| 34 | 1DOI_A | 128 | α/β | 4.67 | 0.570 | 7.35 | 0.534 | 8.53 | 0.397 |
| 35 | 1DP7_P | 76 | α/β | 2.29 | 0.705 | 2.53 | 0.668 | 3.03 | 0.649 |
| 36 | 1DTP_A | 190 | α/β | 18.57 | 0.392 | 19.08 | 0.345 | 17.80 | 0.223 |
| 37 | 1DTV_A | 67 | α/β | 8.36 | 0.302 | 13.88 | 0.205 | 8.96 | 0.250 |
| 38 | 1DUN_A | 120 | α/β | 6.68 | 0.719 | 6.98 | 0.690 | 8.01 | 0.382 |
| 39 | 1DWM_A | 69 | α/β | 3.59 | 0.660 | 4.26 | 0.630 | 3.86 | 0.639 |
| 40 | 1E3Y_A | 104 | α | 3.93 | 0.706 | 4.74 | 0.682 | 8.09 | 0.652 |
| 41 | 1E53_A | 59 | α/β | 3.51 | 0.547 | 3.64 | 0.523 | 4.01 | 0.464 |
| 42 | 1EGG_B | 144 | α/β | 10.48 | 0.683 | 10.56 | 0.677 | 9.19 | 0.433 |
| 43 | 1EKZ_A | 76 | α/β | 6.02 | 0.673 | 6.33 | 0.648 | 4.53 | 0.654 |

| NO. | PDB | Length | Type | GDDfold | | lbfgs-fold | | Rosetta-dist | |
|---|---|---|---|---|---|---|---|---|---|
| | | | | RMSD | TMscore | RMSD | TMscore | RMSD | TMscore |
| 44 | 1ELW_A | 117 | α | 2.43 | 0.784 | 2.61 | 0.761 | 1.37 | 0.907 |
| 45 | 1EM8_D | 112 | α/β | 3.73 | 0.715 | 4.41 | 0.700 | 5.63 | 0.491 |
| 46 | 1EZV_G | 81 | α/β | 12.11 | 0.348 | 12.12 | 0.312 | 9.51 | 0.414 |
| 47 | 1F15_C | 191 | α/β | 25.92 | 0.235 | 25.70 | 0.181 | 21.12 | 0.221 |
| 48 | 1F1E_A | 151 | α/β | 7.19 | 0.590 | 8.47 | 0.466 | 8.99 | 0.492 |
| 49 | 1F2R_I | 100 | α/β | 7.27 | 0.604 | 8.80 | 0.572 | 4.58 | 0.593 |
| 50 | 1F3U_A | 118 | α/β | 20.89 | 0.166 | 23.47 | 0.132 | 17.73 | 0.194 |
| 51 | 1F3Y_A | 165 | α/β | 6.07 | 0.737 | 8.87 | 0.692 | 13.81 | 0.400 |
| 52 | 1F43_A | 61 | α | 4.39 | 0.581 | 7.80 | 0.560 | 4.31 | 0.563 |
| 53 | 1F93_A | 103 | α/β | 2.08 | 0.801 | 3.83 | 0.757 | 2.81 | 0.716 |
| 54 | 1F98_A | 125 | α/β | 7.72 | 0.722 | 9.21 | 0.707 | 8.54 | 0.593 |
| 55 | 1F9P_A | 81 | α/β | 7.61 | 0.600 | 9.23 | 0.568 | 5.25 | 0.576 |
| 56 | 1FAQ_A | 52 | β | 2.73 | 0.601 | 3.25 | 0.555 | 4.69 | 0.446 |
| 57 | 1FC3_A | 116 | α | 2.76 | 0.752 | 2.96 | 0.750 | 3.37 | 0.680 |
| 58 | 1FCA_A | 55 | α/β | 1.43 | 0.783 | 1.59 | 0.742 | 1.86 | 0.699 |
| 59 | 1FEX_A | 59 | α | 2.45 | 0.620 | 2.51 | 0.602 | 2.87 | 0.633 |
| 60 | 1FHT_A | 116 | α/β | 8.53 | 0.689 | 6.97 | 0.679 | 7.89 | 0.518 |
| 61 | 1FJG_F | 101 | α/β | 3.38 | 0.809 | 3.94 | 0.779 | 5.29 | 0.604 |
| 62 | 1FJG_H | 138 | α/β | 2.67 | 0.811 | 3.57 | 0.770 | 11.16 | 0.614 |
| 63 | 1FMB_A | 104 | α/β | 5.16 | 0.721 | 5.36 | 0.679 | 7.09 | 0.443 |
| 64 | 1FR0_A | 125 | α | 4.17 | 0.671 | 4.12 | 0.659 | 3.20 | 0.721 |
| 65 | 1FRD_A | 98 | α/β | 2.48 | 0.748 | 2.68 | 0.735 | 4.85 | 0.535 |
| 66 | 1FSP_A | 124 | α/β | 3.40 | 0.819 | 3.24 | 0.815 | 3.70 | 0.735 |
| 67 | 1FW9_A | 164 | α/β | 3.32 | 0.758 | 8.73 | 0.730 | 9.55 | 0.447 |
| 68 | 1G2R_A | 94 | α/β | 2.01 | 0.810 | 2.15 | 0.796 | 2.98 | 0.697 |
| 69 | 1GGS_A | 81 | α/β | 6.25 | 0.655 | 8.80 | 0.638 | 6.81 | 0.699 |
| 70 | 1GME_A | 150 | α/β | 9.98 | 0.559 | 16.13 | 0.541 | 14.74 | 0.365 |
| 71 | 1GPQ_B | 128 | α/β | 5.19 | 0.663 | 5.36 | 0.624 | 6.74 | 0.546 |
| 72 | 1GQA_A | 130 | α/β | 2.60 | 0.799 | 2.76 | 0.766 | 6.75 | 0.722 |
| 73 | 1GVN_A | 87 | α | 17.00 | 0.235 | 16.57 | 0.219 | 12.42 | 0.317 |
| 74 | 1GVP_A | 87 | α/β | 5.36 | 0.689 | 5.53 | 0.669 | 4.00 | 0.604 |
| 75 | 1GXD_C | 192 | α/β | 13.02 | 0.530 | 14.97 | 0.509 | 14.85 | 0.262 |
| 76 | 1H8E_H | 89 | α/β | 1.66 | 0.840 | 1.85 | 0.825 | 11.57 | 0.527 |
| 77 | 1H9E_A | 56 | α | 4.11 | 0.519 | 4.97 | 0.484 | 5.15 | 0.528 |
| 78 | 1H9F_A | 57 | α | 4.30 | 0.516 | 5.98 | 0.496 | 4.00 | 0.542 |
| 79 | 1HBG_A | 147 | α | 2.09 | 0.855 | 2.08 | 0.852 | 2.80 | 0.799 |
| 80 | 1HBX_E | 89 | α/β | 16.43 | 0.509 | 17.60 | 0.510 | 6.18 | 0.557 |
| 81 | 1HCD_A | 118 | α/β | 3.20 | 0.718 | 3.52 | 0.692 | 6.44 | 0.430 |
| 82 | 1HH8_A | 192 | α/β | 12.60 | 0.695 | 18.87 | 0.643 | 12.11 | 0.690 |
| 83 | 1HHV_A | 74 | α/β | 5.72 | 0.590 | 10.52 | 0.575 | 6.65 | 0.507 |
| 84 | 1HKQ_A | 125 | α/β | 8.85 | 0.685 | 6.76 | 0.674 | 3.65 | 0.637 |
| 85 | 1HKX_E | 143 | α/β | 4.46 | 0.696 | 5.61 | 0.679 | 10.22 | 0.509 |
| 86 | 1HL6_D | 143 | α/β | 4.55 | 0.660 | 5.80 | 0.548 | 6.86 | 0.479 |
| 87 | 1I35_A | 95 | α/β | 3.86 | 0.582 | 4.50 | 0.544 | 5.09 | 0.525 |

| NO. | PDB | Length | Type | GDDfold | | lbfgs-fold | | Rosetta-dist | |
|---|---|---|---|---|---|---|---|---|---|
| | | | | RMSD | TMscore | RMSD | TMscore | RMSD | TMscore |
| 88 | 1I85_A | 110 | β | 2.22 | 0.789 | 2.45 | 0.765 | 5.91 | 0.542 |
| 89 | 1ID2_A | 106 | α/β | 5.60 | 0.738 | 6.78 | 0.695 | 7.23 | 0.535 |
| 90 | 1IM3_D | 95 | α/β | 12.28 | 0.351 | 10.20 | 0.298 | 11.28 | 0.303 |
| 91 | 1IOO_A | 196 | α/β | 4.02 | 0.740 | 5.40 | 0.703 | 11.64 | 0.336 |
| 92 | 1IRS_A | 112 | α/β | 2.62 | 0.811 | 2.96 | 0.792 | 5.10 | 0.554 |
| 93 | 1IS7_K | 85 | α/β | 5.45 | 0.622 | 6.22 | 0.605 | 7.65 | 0.484 |
| 94 | 1IUJ_B | 103 | α/β | 1.89 | 0.837 | 2.42 | 0.818 | 3.65 | 0.687 |
| 95 | 1IUY_A | 92 | α/β | 8.78 | 0.665 | 9.85 | 0.669 | 9.63 | 0.480 |
| 96 | 1J8I_A | 93 | α/β | 9.47 | 0.537 | 10.46 | 0.482 | 12.15 | 0.556 |
| 97 | 1J9I_A | 68 | α/β | 5.41 | 0.556 | 5.66 | 0.528 | 5.92 | 0.582 |
| 98 | 1JEI_A | 53 | α | 6.35 | 0.590 | 4.47 | 0.574 | 6.07 | 0.600 |
| 99 | 1JIW_I | 105 | α/β | 3.03 | 0.732 | 3.54 | 0.696 | 5.87 | 0.489 |
| 100 | 1JJ2_S | 119 | α/β | 4.53 | 0.698 | 5.08 | 0.641 | 5.52 | 0.540 |
| 101 | 1JLI_A | 112 | α | 12.71 | 0.355 | 13.84 | 0.311 | 14.08 | 0.281 |
| 102 | 1JMT_A | 98 | α/β | 4.64 | 0.731 | 4.44 | 0.707 | 4.99 | 0.584 |
| 103 | 1JO0_A | 97 | α/β | 2.05 | 0.826 | 2.94 | 0.798 | 2.44 | 0.742 |
| 104 | 1JOP_A | 140 | α/β | 3.13 | 0.741 | 3.28 | 0.722 | 10.34 | 0.354 |
| 105 | 1JPY_Y | 120 | α/β | 7.65 | 0.578 | 15.24 | 0.531 | 11.22 | 0.365 |
| 106 | 1JR5_A | 90 | α | 7.75 | 0.549 | 7.92 | 0.501 | 8.48 | 0.477 |
| 107 | 1K1Z_A | 78 | β | 3.97 | 0.589 | 6.26 | 0.536 | 5.97 | 0.481 |
| 108 | 1K3S_A | 109 | α/β | 3.23 | 0.689 | 3.32 | 0.684 | 4.97 | 0.538 |
| 109 | 1K5D_B | 146 | α/β | 9.67 | 0.753 | 11.87 | 0.736 | 8.66 | 0.476 |
| 110 | 1K73_1 | 73 | α/β | 5.56 | 0.539 | 6.68 | 0.471 | 3.04 | 0.616 |
| 111 | 1KA8_A | 100 | α/β | 6.31 | 0.570 | 5.75 | 0.561 | 6.98 | 0.546 |
| 112 | 1KN6_A | 73 | α/β | 3.33 | 0.615 | 3.54 | 0.577 | 3.59 | 0.554 |
| 113 | 1KOH_D | 172 | α/β | 6.63 | 0.751 | 6.60 | 0.739 | 8.53 | 0.713 |
| 114 | 1KP6_A | 79 | α/β | 13.25 | 0.285 | 12.89 | 0.267 | 12.36 | 0.234 |
| 115 | 1KPT_A | 105 | α/β | 3.30 | 0.689 | 3.36 | 0.671 | 5.12 | 0.483 |
| 116 | 1KQ6_A | 140 | α/β | 7.43 | 0.706 | 9.08 | 0.667 | 6.76 | 0.536 |
| 117 | 1KSX_A | 144 | α/β | 2.52 | 0.800 | 2.76 | 0.782 | 5.47 | 0.536 |
| 118 | 1KX5_D | 107 | α | 5.34 | 0.569 | 9.89 | 0.507 | 4.08 | 0.817 |
| 119 | 1L1D_B | 147 | α/β | 2.59 | 0.825 | 2.64 | 0.813 | 8.05 | 0.402 |
| 120 | 1L3G_A | 123 | α/β | 6.26 | 0.561 | 10.37 | 0.498 | 6.39 | 0.559 |
| 121 | 1L6H_A | 69 | α | 3.66 | 0.511 | 4.28 | 0.479 | 6.76 | 0.495 |
| 122 | 1LDD_A | 71 | α/β | 1.50 | 0.830 | 1.62 | 0.817 | 2.60 | 0.702 |
| 123 | 1LE2_A | 144 | α | 54.28 | 0.240 | 50.20 | 0.225 | 53.37 | 0.256 |
| 124 | 1LFU_P | 82 | α | 11.52 | 0.554 | 11.90 | 0.529 | 11.79 | 0.589 |
| 125 | 1LNW_C | 139 | α/β | 6.87 | 0.712 | 6.37 | 0.700 | 3.78 | 0.763 |
| 126 | 1LQM_H | 84 | α/β | 5.40 | 0.546 | 8.74 | 0.436 | 6.00 | 0.474 |
| 127 | 1LR1_B | 57 | α | 5.55 | 0.364 | 5.65 | 0.354 | 4.04 | 0.515 |
| 128 | 1LZW_B | 146 | α/β | 2.38 | 0.843 | 2.72 | 0.827 | 6.40 | 0.682 |
| 129 | 1MAI_A | 119 | α/β | 2.27 | 0.821 | 3.37 | 0.773 | 4.90 | 0.518 |
| 130 | 1MFQ_C | 108 | α | 9.60 | 0.588 | 9.65 | 0.578 | 9.11 | 0.566 |
| 131 | 1MWQ_A | 99 | α/β | 2.55 | 0.768 | 2.91 | 0.750 | 3.66 | 0.607 |

| NO. | PDB | Length | Type | GDDfold | | lbfgs-fold | | Rosetta-dist | |
|---|---|---|---|---|---|---|---|---|---|
| | | | | RMSD | TMscore | RMSD | TMscore | RMSD | TMscore |
| 132 | 1N12_A | 138 | α/β | 3.44 | 0.738 | 3.62 | 0.720 | 9.76 | 0.341 |
| 133 | 1N3G_A | 113 | α/β | 7.16 | 0.727 | 7.39 | 0.709 | 6.68 | 0.629 |
| 134 | 1NF6_F | 171 | α/β | 3.03 | 0.774 | 6.29 | 0.742 | 11.07 | 0.709 |
| 135 | 1NGL_A | 179 | α/β | 7.01 | 0.662 | 9.20 | 0.653 | 8.59 | 0.470 |
| 136 | 1NKZ_A | 53 | α | 3.48 | 0.557 | 7.33 | 0.435 | 4.77 | 0.592 |
| 137 | 1NLQ_A | 105 | α/β | 2.34 | 0.803 | 2.84 | 0.767 | 6.31 | 0.400 |
| 138 | 1NOE_A | 86 | α/β | 4.44 | 0.605 | 4.47 | 0.599 | 5.29 | 0.430 |
| 139 | 1NPB_A | 140 | α/β | 7.82 | 0.678 | 8.38 | 0.642 | 8.01 | 0.589 |
| 140 | 1NQZ_A | 171 | α/β | 6.20 | 0.784 | 5.62 | 0.774 | 11.09 | 0.381 |
| 141 | 1NR3_A | 122 | α/β | 17.46 | 0.319 | 14.59 | 0.308 | 11.61 | 0.383 |
| 142 | 1NTV_A | 152 | α/β | 4.01 | 0.758 | 4.29 | 0.747 | 6.15 | 0.477 |
| 143 | 1NZE_A | 112 | α | 3.58 | 0.839 | 3.13 | 0.844 | 4.29 | 0.813 |
| 144 | 1O0G_A | 124 | α/β | 4.78 | 0.646 | 5.60 | 0.617 | 6.73 | 0.443 |
| 145 | 1OA8_D | 133 | α/β | 13.11 | 0.369 | 15.06 | 0.295 | 14.96 | 0.264 |
| 146 | 1OFT_A | 119 | α/β | 2.52 | 0.791 | 2.60 | 0.783 | 4.43 | 0.575 |
| 147 | 1OJG_A | 136 | α/β | 3.93 | 0.635 | 3.98 | 0.625 | 5.27 | 0.522 |
| 148 | 1OOF_A | 124 | α/β | 2.11 | 0.830 | 2.42 | 0.804 | 6.21 | 0.672 |
| 149 | 1ORY_A | 119 | α | 3.58 | 0.756 | 3.84 | 0.750 | 3.90 | 0.678 |
| 150 | 1OX7_A | 158 | α/β | 3.08 | 0.795 | 4.13 | 0.780 | 5.24 | 0.565 |
| 151 | 1OZ9_A | 141 | α/β | 2.62 | 0.792 | 4.15 | 0.748 | 6.62 | 0.578 |
| 152 | 1PD6_A | 94 | β | 5.14 | 0.629 | 5.00 | 0.601 | 4.56 | 0.644 |
| 153 | 1PFS_A | 78 | β | 4.97 | 0.573 | 5.49 | 0.553 | 5.31 | 0.433 |
| 154 | 1PGV_A | 167 | α/β | 3.23 | 0.773 | 3.90 | 0.729 | 3.69 | 0.705 |
| 155 | 1PIH_A | 73 | α/β | 4.53 | 0.620 | 6.06 | 0.585 | 6.97 | 0.403 |
| 156 | 1PMS_A | 135 | α/β | 5.83 | 0.632 | 8.51 | 0.596 | 10.86 | 0.520 |
| 157 | 1PSR_A | 100 | α/β | 4.80 | 0.677 | 6.81 | 0.686 | 3.86 | 0.635 |
| 158 | 1PXW_A | 128 | α/β | 5.57 | 0.717 | 6.95 | 0.709 | 5.53 | 0.534 |
| 159 | 1PZW_A | 80 | α/β | 2.52 | 0.731 | 3.07 | 0.684 | 4.20 | 0.519 |
| 160 | 1QFT_A | 175 | α/β | 3.94 | 0.773 | 4.52 | 0.768 | 8.21 | 0.460 |
| 161 | 1QFW_B | 110 | β | 7.77 | 0.576 | 9.34 | 0.488 | 8.64 | 0.347 |
| 162 | 1QMA_A | 123 | α/β | 1.74 | 0.877 | 1.80 | 0.872 | 4.71 | 0.562 |
| 163 | 1QZG_A | 170 | α/β | 5.15 | 0.700 | 5.24 | 0.661 | 9.04 | 0.419 |
| 164 | 1R5T_A | 141 | α/β | 3.10 | 0.810 | 2.88 | 0.798 | 5.42 | 0.586 |
| 165 | 1R6R_A | 80 | α | 8.42 | 0.460 | 7.61 | 0.444 | 8.48 | 0.556 |
| 166 | 1RHX_A | 87 | α/β | 4.28 | 0.615 | 4.37 | 0.603 | 4.63 | 0.486 |
| 167 | 1ROW_A | 107 | α/β | 2.03 | 0.819 | 2.15 | 0.802 | 4.71 | 0.495 |
| 168 | 1RTU_A | 114 | α/β | 3.48 | 0.696 | 3.77 | 0.677 | 6.98 | 0.529 |
| 169 | 1RZ3_A | 184 | α/β | 3.52 | 0.817 | 5.14 | 0.775 | 8.85 | 0.511 |
| 170 | 1S2D_A | 165 | α/β | 5.30 | 0.763 | 6.55 | 0.709 | 7.62 | 0.542 |
| 171 | 1S3J_A | 143 | α/β | 4.83 | 0.638 | 4.87 | 0.636 | 2.89 | 0.753 |
| 172 | 1S56_B | 135 | α/β | 6.68 | 0.754 | 7.27 | 0.739 | 7.27 | 0.670 |
| 173 | 1S7O_C | 108 | α | 8.07 | 0.540 | 7.33 | 0.530 | 10.47 | 0.557 |
| 174 | 1S7Z_A | 106 | α | 4.83 | 0.629 | 6.18 | 0.607 | 6.80 | 0.572 |
| 175 | 1SAU_A | 114 | α/β | 3.01 | 0.709 | 3.23 | 0.686 | 4.30 | 0.643 |

| NO. | PDB | Length | Type | GDDfold | | lbfgs-fold | | Rosetta-dist | |
|---|---|---|---|---|---|---|---|---|---|
| | | | | RMSD | TMscore | RMSD | TMscore | RMSD | TMscore |
| 176 | 1SMP_I | 100 | α/β | 2.53 | 0.764 | 2.40 | 0.772 | 5.73 | 0.532 |
| 177 | 1SPP_B | 112 | α/β | 2.70 | 0.762 | 5.18 | 0.711 | 5.62 | 0.432 |
| 178 | 1STM_A | 141 | α/β | 5.35 | 0.594 | 7.34 | 0.533 | 12.64 | 0.274 |
| 179 | 1SVJ_A | 136 | α/β | 2.58 | 0.787 | 2.79 | 0.775 | 6.59 | 0.472 |
| 180 | 1TEO_A | 173 | α/β | 4.96 | 0.718 | 7.11 | 0.694 | 7.99 | 0.509 |
| 181 | 1TJF_B | 186 | α/β | 3.98 | 0.837 | 5.87 | 0.814 | 5.44 | 0.722 |
| 182 | 1TLJ_A | 189 | α/β | 3.45 | 0.778 | 3.87 | 0.738 | 7.31 | 0.492 |
| 183 | 1TUL_A | 102 | α/β | 3.98 | 0.666 | 3.99 | 0.626 | 8.48 | 0.341 |
| 184 | 1TWU_A | 137 | α/β | 4.13 | 0.716 | 4.46 | 0.691 | 3.95 | 0.689 |
| 185 | 1TYG_B | 65 | α/β | 2.06 | 0.755 | 2.45 | 0.745 | 2.68 | 0.672 |
| 186 | 1TZ0_A | 108 | α/β | 3.18 | 0.717 | 4.90 | 0.661 | 5.66 | 0.489 |
| 187 | 1U84_A | 81 | α | 1.89 | 0.809 | 1.99 | 0.801 | 1.93 | 0.825 |
| 188 | 1UFB_A | 127 | α | 2.38 | 0.817 | 3.25 | 0.784 | 4.77 | 0.683 |
| 189 | 1UG4_A | 60 | β | 5.74 | 0.623 | 4.07 | 0.600 | 5.41 | 0.379 |
| 190 | 1UNG_D | 149 | α | 4.17 | 0.705 | 4.52 | 0.693 | 4.55 | 0.640 |
| 191 | 1USL_C | 158 | α/β | 3.93 | 0.851 | 5.59 | 0.843 | 5.23 | 0.626 |
| 192 | 1V74_A | 107 | α/β | 6.42 | 0.639 | 4.58 | 0.637 | 4.88 | 0.623 |
| 193 | 1VCC_A | 77 | α/β | 5.59 | 0.509 | 7.79 | 0.450 | 4.22 | 0.563 |
| 194 | 1VCY_A | 193 | α/β | 4.29 | 0.735 | 6.57 | 0.712 | 9.12 | 0.394 |
| 195 | 1VD0_A | 109 | α/β | 12.54 | 0.651 | 12.91 | 0.627 | 10.75 | 0.333 |
| 196 | 1VHG_A | 185 | α/β | 12.70 | 0.665 | 14.03 | 0.651 | 10.51 | 0.338 |
| 197 | 1VKE_E | 119 | α | 10.91 | 0.643 | 11.01 | 0.627 | 5.66 | 0.662 |
| 198 | 1VYI_A | 111 | α/β | 21.15 | 0.232 | 18.30 | 0.256 | 17.01 | 0.263 |
| 199 | 1VYX_A | 60 | α/β | 5.78 | 0.457 | 6.03 | 0.427 | 4.71 | 0.371 |
| 200 | 1W1W_E | 70 | α/β | 2.25 | 0.782 | 2.25 | 0.771 | 2.56 | 0.712 |
| 201 | 1WJ8_A | 117 | α | 1.79 | 0.860 | 1.96 | 0.847 | 4.56 | 0.758 |
| 202 | 1WLQ_C | 185 | α/β | 6.92 | 0.640 | 7.19 | 0.598 | 9.86 | 0.460 |
| 203 | 1WMH_B | 82 | α/β | 1.70 | 0.832 | 1.85 | 0.807 | 3.63 | 0.561 |
| 204 | 1XJA_C | 169 | α/β | 4.15 | 0.748 | 5.81 | 0.719 | 4.95 | 0.622 |
| 205 | 1Y14_A | 133 | α | 9.56 | 0.525 | 12.03 | 0.413 | 7.23 | 0.578 |
| 206 | 1Y1X_A | 182 | α/β | 5.23 | 0.653 | 6.74 | 0.628 | 5.75 | 0.624 |
| 207 | 1YG2_A | 169 | α/β | 11.50 | 0.483 | 20.26 | 0.446 | 5.25 | 0.551 |
| 208 | 1Z8R_A | 150 | α/β | 12.46 | 0.471 | 13.75 | 0.406 | 12.34 | 0.298 |
| 209 | 2A5Y_A | 173 | α | 3.95 | 0.744 | 5.46 | 0.690 | 4.91 | 0.645 |
| 210 | 2A9U_B | 127 | α | 11.29 | 0.658 | 10.80 | 0.652 | 9.22 | 0.754 |
| 211 | 2ACY_A | 98 | α/β | 3.16 | 0.793 | 3.13 | 0.761 | 3.65 | 0.629 |
| 212 | 2AEN_A | 164 | α/β | 6.27 | 0.621 | 10.17 | 0.496 | 11.68 | 0.323 |
| 213 | 2APN_A | 114 | α/β | 7.98 | 0.651 | 7.95 | 0.619 | 6.89 | 0.492 |
| 214 | 2AQ0_A | 84 | α | 7.10 | 0.637 | 13.47 | 0.576 | 7.07 | 0.644 |
| 215 | 2AQS_A | 160 | α/β | 4.11 | 0.766 | 4.18 | 0.737 | 11.18 | 0.388 |
| 216 | 2BSE_A | 107 | α/β | 4.27 | 0.601 | 5.40 | 0.503 | 8.37 | 0.423 |
| 217 | 2BWJ_A | 196 | α/β | 4.06 | 0.780 | 5.91 | 0.758 | 7.29 | 0.526 |
| 218 | 2BYK_D | 92 | α | 4.42 | 0.586 | 5.88 | 0.543 | 1.82 | 0.852 |
| 219 | 2C2F_A | 178 | α/β | 3.76 | 0.807 | 4.77 | 0.804 | 6.81 | 0.693 |

| NO. | PDB | Length | Type | GDDfold | | lbfgs-fold | | Rosetta-dist | |
|---|---|---|---|---|---|---|---|---|---|
| | | | | RMSD | TMscore | RMSD | TMscore | RMSD | TMscore |
| 220 | 2C4W_A | 168 | α/β | 2.84 | 0.831 | 3.97 | 0.816 | 6.28 | 0.505 |
| 221 | 2CDP_A | 138 | α/β | 3.07 | 0.784 | 3.07 | 0.769 | 6.85 | 0.461 |
| 222 | 2CMX_A | 70 | α/β | 2.06 | 0.740 | 2.58 | 0.701 | 3.19 | 0.649 |
| 223 | 2CO3_B | 135 | α/β | 6.35 | 0.670 | 6.40 | 0.642 | 10.63 | 0.315 |
| 224 | 2CWP_A | 109 | α/β | 5.73 | 0.733 | 6.08 | 0.720 | 6.13 | 0.478 |
| 225 | 2CZV_D | 119 | α/β | 2.37 | 0.816 | 5.94 | 0.765 | 5.49 | 0.620 |
| 226 | 2D0P_B | 110 | α/β | 2.94 | 0.794 | 2.91 | 0.781 | 8.77 | 0.638 |
| 227 | 2EWC_B | 122 | α/β | 3.42 | 0.765 | 4.21 | 0.747 | 7.50 | 0.637 |
| 228 | 2F22_B | 143 | α/β | 3.14 | 0.789 | 3.35 | 0.774 | 5.67 | 0.563 |
| 229 | 2FA5_B | 142 | α/β | 7.84 | 0.723 | 7.14 | 0.690 | 4.54 | 0.708 |
| 230 | 2FKB_C | 167 | α/β | 2.96 | 0.804 | 4.48 | 0.796 | 8.73 | 0.425 |
| 231 | 2GBJ_B | 84 | α/β | 2.58 | 0.756 | 2.33 | 0.764 | 4.10 | 0.627 |
| 232 | 2GJ3_A | 119 | α/β | 2.70 | 0.805 | 2.90 | 0.804 | 4.79 | 0.633 |
| 233 | 2GKC_A | 155 | α/β | 4.97 | 0.629 | 4.84 | 0.623 | 6.72 | 0.421 |
| 234 | 2H30_A | 151 | α/β | 4.73 | 0.702 | 5.94 | 0.687 | 6.96 | 0.549 |
| 235 | 2H8E_A | 120 | α/β | 2.80 | 0.770 | 2.79 | 0.769 | 5.50 | 0.506 |
| 236 | 2HI3_A | 73 | α | 5.70 | 0.616 | 5.24 | 0.612 | 4.73 | 0.637 |
| 237 | 2HQ7_B | 142 | α/β | 2.98 | 0.797 | 4.25 | 0.751 | 5.71 | 0.535 |
| 238 | 2HYB_A | 130 | α/β | 2.15 | 0.830 | 2.57 | 0.810 | 5.09 | 0.548 |
| 239 | 2ICT_A | 94 | α | 7.48 | 0.729 | 8.32 | 0.716 | 3.91 | 0.765 |
| 240 | 2J4H_B | 172 | α/β | 7.55 | 0.727 | 7.59 | 0.716 | 12.24 | 0.374 |
| 241 | 2J6Z_A | 86 | α | 3.72 | 0.640 | 4.13 | 0.607 | 2.50 | 0.733 |
| 242 | 2JLP_D | 169 | α/β | 5.30 | 0.710 | 5.60 | 0.653 | 10.19 | 0.360 |
| 243 | 2JP3_A | 67 | α | 10.11 | 0.307 | 10.26 | 0.296 | 6.04 | 0.362 |
| 244 | 2K9X_A | 102 | α/β | 4.53 | 0.677 | 4.36 | 0.655 | 5.80 | 0.469 |
| 245 | 2KBW_A | 160 | α | 3.45 | 0.742 | 4.38 | 0.729 | 4.28 | 0.715 |
| 246 | 2L5P_A | 175 | α/β | 4.23 | 0.758 | 3.94 | 0.767 | 6.61 | 0.515 |
| 247 | 2L74_A | 125 | α/β | 6.95 | 0.702 | 8.79 | 0.684 | 5.35 | 0.546 |
| 248 | 2LKP_A | 119 | α/β | 9.48 | 0.602 | 10.24 | 0.590 | 9.59 | 0.610 |
| 249 | 2LRB_A | 165 | α/β | 3.95 | 0.720 | 4.71 | 0.697 | 5.77 | 0.543 |
| 250 | 2LWP_A | 97 | α/β | 4.90 | 0.550 | 10.34 | 0.539 | 6.75 | 0.483 |
| 251 | 2NAZ_A | 109 | α/β | 4.62 | 0.662 | 4.89 | 0.639 | 4.09 | 0.649 |
| 252 | 2NCM_A | 99 | β | 1.77 | 0.834 | 1.96 | 0.816 | 4.24 | 0.594 |
| 253 | 2NDP_A | 99 | α/β | 8.44 | 0.508 | 8.41 | 0.412 | 5.81 | 0.503 |
| 254 | 2NS9_B | 152 | α/β | 4.02 | 0.814 | 3.94 | 0.815 | 6.51 | 0.525 |
| 255 | 2O70_F | 168 | α | 2.74 | 0.795 | 2.76 | 0.790 | 5.94 | 0.551 |
| 256 | 2ODM_B | 83 | α | 2.93 | 0.709 | 3.26 | 0.699 | 1.85 | 0.852 |
| 257 | 2P7L_A | 125 | α/β | 4.17 | 0.749 | 4.73 | 0.717 | 4.23 | 0.637 |
| 258 | 2PI2_F | 119 | α/β | 3.26 | 0.761 | 5.72 | 0.737 | 5.18 | 0.539 |
| 259 | 2PYB_A | 151 | α | 3.39 | 0.817 | 3.50 | 0.803 | 4.51 | 0.690 |
| 260 | 2Q2H_A | 118 | α/β | 9.37 | 0.637 | 8.85 | 0.623 | 9.66 | 0.378 |
| 261 | 2QVG_A | 129 | α/β | 2.23 | 0.819 | 2.35 | 0.815 | 3.64 | 0.713 |
| 262 | 2QZJ_A | 121 | α/β | 1.54 | 0.889 | 1.61 | 0.882 | 4.31 | 0.751 |
| 263 | 2RD5_D | 126 | α/β | 10.74 | 0.646 | 14.40 | 0.569 | 9.89 | 0.438 |

| NO. | PDB | Length | Type | GDDfold | | lbfgs-fold | | Rosetta-dist | |
|---|---|---|---|---|---|---|---|---|---|
| | | | | RMSD | TMscore | RMSD | TMscore | RMSD | TMscore |
| 264 | 2RLD_C | 116 | α | 2.39 | 0.824 | 2.55 | 0.821 | 2.85 | 0.783 |
| 265 | 2UUX_A | 55 | α/β | 3.60 | 0.649 | 6.56 | 0.615 | 4.71 | 0.387 |
| 266 | 2V85_A | 74 | α/β | 9.08 | 0.379 | 11.78 | 0.245 | 5.97 | 0.460 |
| 267 | 2VUL_A | 193 | α/β | 2.54 | 0.830 | 2.79 | 0.807 | 11.73 | 0.315 |
| 268 | 2WCW_B | 122 | α/β | 3.25 | 0.796 | 3.35 | 0.774 | 4.38 | 0.623 |
| 269 | 2WGP_A | 168 | α/β | 2.85 | 0.809 | 5.96 | 0.776 | 4.98 | 0.662 |
| 270 | 2XGY_A | 129 | α | 5.69 | 0.684 | 7.54 | 0.615 | 6.96 | 0.627 |
| 271 | 2Z3B_A | 180 | α/β | 2.44 | 0.842 | 3.60 | 0.814 | 13.56 | 0.309 |
| 272 | 2ZMZ_B | 79 | α/β | 4.58 | 0.697 | 3.86 | 0.676 | 4.95 | 0.568 |
| 273 | 3ALU_A | 157 | α/β | 3.85 | 0.733 | 6.11 | 0.689 | 8.82 | 0.392 |
| 274 | 3BDB_A | 126 | α/β | 6.60 | 0.733 | 7.93 | 0.712 | 5.78 | 0.499 |
| 275 | 3CAE_A | 132 | α/β | 27.29 | 0.504 | 27.14 | 0.459 | 22.71 | 0.426 |
| 276 | 3CG4_A | 126 | α/β | 2.69 | 0.822 | 2.79 | 0.814 | 3.72 | 0.743 |
| 277 | 3CX5_F | 74 | α | 2.92 | 0.637 | 3.74 | 0.588 | 3.25 | 0.632 |
| 278 | 3CX5_G | 126 | α | 7.28 | 0.640 | 7.52 | 0.608 | 6.43 | 0.466 |
| 279 | 3E6M_E | 147 | α/β | 7.95 | 0.709 | 8.18 | 0.678 | 9.01 | 0.625 |
| 280 | 3E9T_D | 102 | β | 1.77 | 0.849 | 2.04 | 0.811 | 4.45 | 0.546 |
| 281 | 3EOD_A | 115 | α/β | 2.16 | 0.834 | 4.08 | 0.777 | 4.44 | 0.721 |
| 282 | 3F8L_A | 162 | α/β | 3.03 | 0.814 | 4.14 | 0.794 | 9.19 | 0.433 |
| 283 | 3G20_B | 119 | α/β | 7.46 | 0.642 | 7.59 | 0.606 | 5.82 | 0.557 |
| 284 | 3GMX_A | 153 | α/β | 14.33 | 0.410 | 17.32 | 0.398 | 8.09 | 0.357 |
| 285 | 3H05_B | 163 | α/β | 3.68 | 0.801 | 4.30 | 0.778 | 7.27 | 0.473 |
| 286 | 3I9V_7 | 127 | α/β | 5.20 | 0.626 | 5.36 | 0.606 | 6.11 | 0.493 |
| 287 | 3IAM_2 | 179 | α/β | 4.23 | 0.715 | 11.49 | 0.458 | 6.12 | 0.548 |
| 288 | 3LQV_B | 115 | α/β | 5.20 | 0.675 | 9.26 | 0.645 | 8.01 | 0.524 |
| 289 | 3M1N_B | 168 | α/β | 9.63 | 0.510 | 19.92 | 0.446 | 14.27 | 0.391 |
| 290 | 3MQK_C | 75 | β | 1.74 | 0.800 | 1.75 | 0.798 | 3.36 | 0.559 |
| 291 | 3N1G_C | 104 | α/β | 2.18 | 0.818 | 3.00 | 0.791 | 4.06 | 0.611 |
| 292 | 3N9U_C | 96 | α/β | 5.84 | 0.737 | 6.85 | 0.733 | 6.56 | 0.576 |
| 293 | 3O61_A | 187 | α/β | 13.05 | 0.690 | 13.51 | 0.688 | 14.61 | 0.347 |
| 294 | 3P8B_A | 60 | α/β | 2.17 | 0.663 | 3.56 | 0.570 | 4.69 | 0.356 |
| 295 | 3PD2_A | 147 | α/β | 3.23 | 0.776 | 3.70 | 0.736 | 7.75 | 0.541 |
| 296 | 3QU3_A | 122 | α/β | 4.29 | 0.656 | 5.46 | 0.617 | 7.15 | 0.452 |
| 297 | 3SDL_B | 97 | α | 31.18 | 0.237 | 26.55 | 0.214 | 22.36 | 0.333 |
| 298 | 3UE6_E | 138 | α/β | 5.65 | 0.729 | 5.49 | 0.716 | 7.34 | 0.576 |
| 299 | 3V1O_A | 165 | α/β | 4.25 | 0.762 | 6.42 | 0.727 | 8.35 | 0.413 |
| 300 | 3W1Z_D | 110 | α/β | 3.62 | 0.739 | 11.25 | 0.707 | 6.88 | 0.498 |
| 301 | 3X0G_A | 93 | α | 3.32 | 0.665 | 3.87 | 0.618 | 5.94 | 0.520 |
| 302 | 3X15_A | 87 | α | 15.06 | 0.419 | 18.98 | 0.386 | 9.65 | 0.481 |
| 303 | 4AIH_A | 139 | α/β | 4.22 | 0.745 | 5.02 | 0.725 | 2.81 | 0.763 |
| 304 | 4ASW_C | 81 | α/β | 1.95 | 0.796 | 2.08 | 0.782 | 3.73 | 0.573 |
| 305 | 4B0M_A | 131 | α/β | 4.52 | 0.718 | 5.88 | 0.665 | 6.48 | 0.516 |
| 306 | 4CXT_A | 132 | α/β | 6.22 | 0.728 | 6.33 | 0.711 | 6.02 | 0.573 |
| 307 | 4DYW_A | 129 | α/β | 1.84 | 0.865 | 2.08 | 0.850 | 6.45 | 0.431 |

| NO. | PDB | Length | Type | GDDfold | | lbfgs-fold | | Rosetta-dist | |
|---|---|---|---|---|---|---|---|---|---|
| | | | | RMSD | TMscore | RMSD | TMscore | RMSD | TMscore |
| 308 | 4ESB_A | 103 | α/β | 5.57 | 0.745 | 5.43 | 0.729 | 3.08 | 0.720 |
| 309 | 4GDK_A | 88 | α/β | 3.23 | 0.770 | 2.39 | 0.761 | 4.44 | 0.542 |
| 310 | 4GF3_A | 123 | α/β | 7.13 | 0.696 | 7.17 | 0.683 | 8.23 | 0.545 |
| 311 | 4GQY_A | 147 | α/β | 8.77 | 0.747 | 9.42 | 0.736 | 6.55 | 0.543 |
| 312 | 4I60_A | 128 | α/β | 3.01 | 0.777 | 3.35 | 0.749 | 5.21 | 0.553 |
| 313 | 4IOS_A | 100 | α/β | 2.76 | 0.707 | 2.86 | 0.697 | 6.13 | 0.421 |
| 314 | 4J20_A | 88 | α/β | 2.68 | 0.746 | 2.86 | 0.710 | 4.33 | 0.584 |
| 315 | 4JGX_B | 128 | α/β | 3.76 | 0.718 | 4.33 | 0.700 | 4.91 | 0.671 |
| 316 | 4K1F_A | 198 | α/β | 3.65 | 0.781 | 4.46 | 0.761 | 6.76 | 0.562 |
| 317 | 4KA0_A | 143 | α/β | 2.25 | 0.852 | 2.42 | 0.844 | 4.26 | 0.609 |
| 318 | 4LE0_B | 133 | α/β | 1.88 | 0.874 | 2.03 | 0.866 | 2.48 | 0.776 |
| 319 | 4LMS_A | 80 | α/β | 8.25 | 0.462 | 14.92 | 0.327 | 6.59 | 0.347 |
| 320 | 4M75_F | 75 | α/β | 4.59 | 0.759 | 5.91 | 0.743 | 4.82 | 0.593 |
| 321 | 4MLF_D | 61 | β | 14.00 | 0.286 | 12.40 | 0.217 | 8.91 | 0.253 |
| 322 | 4MMG_A | 91 | α/β | 2.06 | 0.813 | 2.20 | 0.797 | 3.89 | 0.644 |
| 323 | 4NBI_A | 163 | α/β | 3.32 | 0.810 | 3.66 | 0.798 | 8.26 | 0.441 |
| 324 | 4OW1_A | 86 | α/β | 3.05 | 0.713 | 2.97 | 0.698 | 4.21 | 0.581 |
| 325 | 4Q2O_A | 92 | α/β | 5.04 | 0.768 | 5.24 | 0.745 | 4.12 | 0.656 |
| 326 | 4Q2Q_A | 90 | α/β | 2.80 | 0.786 | 3.58 | 0.767 | 5.29 | 0.591 |
| 327 | 4R67_0 | 199 | α/β | 2.63 | 0.851 | 2.94 | 0.831 | 14.75 | 0.327 |
| 328 | 4RUV_A | 106 | α/β | 1.87 | 0.832 | 2.11 | 0.819 | 2.51 | 0.742 |
| 329 | 4UIJ_A | 104 | α/β | 1.95 | 0.814 | 2.29 | 0.783 | 2.72 | 0.725 |
| 330 | 4V2O_A | 78 | α | 3.05 | 0.653 | 3.35 | 0.632 | 1.57 | 0.825 |
| 331 | 4Z6J_A | 133 | β | 3.04 | 0.738 | 3.76 | 0.722 | 7.05 | 0.493 |
| 332 | 4ZBY_A | 194 | α/β | 2.21 | 0.868 | 2.27 | 0.859 | 9.28 | 0.433 |
| 333 | 5CJ3_B | 126 | α/β | 3.42 | 0.789 | 3.64 | 0.770 | 3.79 | 0.708 |
| 334 | 5E4E_A | 111 | α/β | 8.80 | 0.384 | 12.74 | 0.304 | 13.88 | 0.248 |
| 335 | 5EKT_A | 196 | α/β | 3.36 | 0.834 | 3.26 | 0.823 | 8.30 | 0.415 |
| 336 | 5IAO_A | 171 | α/β | 3.75 | 0.773 | 3.87 | 0.757 | 10.22 | 0.426 |
| 337 | 5IZB_A | 89 | α/β | 5.75 | 0.567 | 8.57 | 0.539 | 8.70 | 0.468 |
| 338 | 5JTM_A | 155 | α/β | 10.91 | 0.703 | 11.42 | 0.698 | 10.14 | 0.451 |
| 339 | 5L38_A | 91 | α/β | 2.17 | 0.825 | 2.09 | 0.815 | 10.38 | 0.693 |
| 340 | 5L8R_D | 143 | α/β | 12.72 | 0.560 | 13.55 | 0.536 | 11.22 | 0.361 |
| 341 | 5O2V_A | 92 | α/β | 4.96 | 0.755 | 6.15 | 0.717 | 2.99 | 0.703 |
| 342 | 5O8G_A | 122 | α/β | 3.46 | 0.741 | 5.75 | 0.712 | 6.90 | 0.551 |
| 343 | 5T17_A | 85 | α/β | 3.13 | 0.655 | 3.20 | 0.646 | 2.89 | 0.657 |
| 344 | 5TMF_E | 95 | α/β | 7.64 | 0.545 | 6.39 | 0.524 | 6.87 | 0.514 |
| 345 | 5TUV_B | 104 | α/β | 13.95 | 0.377 | 13.62 | 0.337 | 10.45 | 0.467 |
| 346 | 5WSE_A | 114 | α/β | 3.66 | 0.792 | 4.64 | 0.749 | 4.90 | 0.621 |
| 347 | 6AQ3_B | 171 | α/β | 3.56 | 0.752 | 4.73 | 0.700 | 6.67 | 0.533 |

**Table S3**. Prediction results of GDDfold, GDDfold_global, and GDDfold_local for 347 benchmark proteins.

| NO. | PDB | Length | Type | GDDfold | | GDDfold_global | | GDDfold_local | |
|---|---|---|---|---|---|---|---|---|---|
| | | | | RMSD | TMscore | RMSD | TMscore | RMSD | TMscore |
| 1 | 1A3A_C | 146 | α/β | 2.30 | 0.854 | 8.42 | 0.360 | 2.37 | 0.845 |
| 2 | 1A6L_A | 106 | α/β | 2.57 | 0.749 | 12.08 | 0.379 | 2.70 | 0.740 |
| 3 | 1A7D_A | 118 | α | 4.35 | 0.808 | 10.66 | 0.353 | 3.70 | 0.802 |
| 4 | 1A91_A | 79 | α | 4.95 | 0.461 | 6.10 | 0.368 | 5.06 | 0.455 |
| 5 | 1ABT_A | 74 | α/β | 4.79 | 0.576 | 12.09 | 0.288 | 4.30 | 0.562 |
| 6 | 1ABV_A | 105 | α | 3.04 | 0.698 | 10.12 | 0.348 | 3.14 | 0.684 |
| 7 | 1AHK_A | 129 | β | 5.79 | 0.531 | 16.44 | 0.231 | 6.01 | 0.519 |
| 8 | 1AK6_A | 174 | α/β | 6.03 | 0.670 | 12.10 | 0.363 | 6.07 | 0.646 |
| 9 | 1AKP_A | 114 | β | 3.07 | 0.712 | 18.00 | 0.217 | 3.73 | 0.678 |
| 10 | 1AP7_A | 168 | α | 3.37 | 0.761 | 7.35 | 0.420 | 5.08 | 0.749 |
| 11 | 1AUU_A | 55 | β | 2.12 | 0.680 | 11.03 | 0.343 | 2.43 | 0.667 |
| 12 | 1AX8_A | 130 | α/β | 4.06 | 0.687 | 17.93 | 0.284 | 5.54 | 0.634 |
| 13 | 1B4R_A | 80 | β | 2.33 | 0.743 | 11.86 | 0.252 | 2.48 | 0.722 |
| 14 | 1B4U_A | 132 | α | 11.38 | 0.597 | 13.47 | 0.441 | 7.58 | 0.571 |
| 15 | 1B8Q_A | 127 | α/β | 9.19 | 0.505 | 16.45 | 0.301 | 9.82 | 0.454 |
| 16 | 1BE3_J | 62 | α | 9.63 | 0.413 | 10.39 | 0.320 | 9.31 | 0.372 |
| 17 | 1BGF_A | 124 | α/β | 2.82 | 0.734 | 7.32 | 0.475 | 2.93 | 0.720 |
| 18 | 1BGY_J | 62 | α | 9.45 | 0.381 | 11.27 | 0.329 | 10.23 | 0.362 |
| 19 | 1BJX_A | 110 | α/β | 9.74 | 0.746 | 8.14 | 0.367 | 9.93 | 0.729 |
| 20 | 1BUO_A | 121 | α/β | 2.93 | 0.776 | 13.44 | 0.479 | 7.87 | 0.761 |
| 21 | 1C03_A | 163 | α/β | 11.58 | 0.609 | 17.62 | 0.322 | 12.14 | 0.596 |
| 22 | 1C41_A | 165 | α/β | 3.45 | 0.844 | 14.29 | 0.413 | 4.09 | 0.830 |
| 23 | 1C9F_A | 87 | α/β | 3.47 | 0.612 | 5.99 | 0.394 | 4.39 | 0.589 |
| 24 | 1CDB_A | 105 | β | 4.67 | 0.706 | 10.83 | 0.406 | 4.33 | 0.685 |
| 25 | 1CF7_B | 82 | α/β | 2.57 | 0.715 | 10.52 | 0.300 | 2.84 | 0.660 |
| 26 | 1CHC_A | 68 | α/β | 5.59 | 0.513 | 8.52 | 0.288 | 5.52 | 0.496 |
| 27 | 1CTO_A | 109 | β | 4.87 | 0.701 | 14.98 | 0.262 | 5.27 | 0.657 |
| 28 | 1CXZ_B | 86 | α | 5.22 | 0.621 | 7.73 | 0.327 | 4.69 | 0.603 |
| 29 | 1D6T_A | 117 | α/β | 4.60 | 0.753 | 10.92 | 0.419 | 4.83 | 0.742 |
| 30 | 1D8B_A | 81 | α | 2.46 | 0.715 | 7.87 | 0.330 | 2.53 | 0.704 |
| 31 | 1DBF_A | 127 | α/β | 4.02 | 0.802 | 14.19 | 0.326 | 5.05 | 0.784 |
| 32 | 1DCF_A | 133 | α/β | 3.21 | 0.813 | 4.74 | 0.629 | 4.43 | 0.806 |
| 33 | 1DL6_A | 58 | β | 11.74 | 0.378 | 8.52 | 0.409 | 11.84 | 0.345 |
| 34 | 1DOI_A | 128 | α/β | 4.67 | 0.570 | 17.19 | 0.257 | 7.51 | 0.537 |
| 35 | 1DP7_P | 76 | α/β | 2.29 | 0.705 | 7.13 | 0.412 | 2.37 | 0.691 |
| 36 | 1DTP_A | 190 | α/β | 18.57 | 0.392 | 29.85 | 0.139 | 18.02 | 0.242 |
| 37 | 1DTV_A | 67 | α/β | 8.36 | 0.302 | 21.08 | 0.213 | 8.00 | 0.259 |
| 38 | 1DUN_A | 120 | α/β | 6.68 | 0.719 | 19.24 | 0.256 | 9.89 | 0.683 |
| 39 | 1DWM_A | 69 | α/β | 3.59 | 0.660 | 11.45 | 0.369 | 3.95 | 0.647 |
| 40 | 1E3Y_A | 104 | α | 3.93 | 0.706 | 10.25 | 0.315 | 4.42 | 0.690 |
| 41 | 1E53_A | 59 | α/β | 3.51 | 0.547 | 12.26 | 0.244 | 3.69 | 0.531 |
| 42 | 1EGG_B | 144 | α/β | 10.48 | 0.683 | 16.53 | 0.230 | 10.68 | 0.675 |

| NO. | PDB | Length | Type | GDDfold | | GDDfold_global | | GDDfold_local | |
|---|---|---|---|---|---|---|---|---|---|
| | | | | RMSD | TMscore | RMSD | TMscore | RMSD | TMscore |
| 43 | 1EKZ_A | 76 | α/β | 6.02 | 0.673 | 9.90 | 0.364 | 5.57 | 0.666 |
| 44 | 1ELW_A | 117 | α | 2.43 | 0.784 | 3.40 | 0.661 | 2.52 | 0.773 |
| 45 | 1EM8_D | 112 | α/β | 3.73 | 0.715 | 9.06 | 0.402 | 3.96 | 0.701 |
| 46 | 1EZV_G | 81 | α/β | 12.11 | 0.348 | 14.24 | 0.329 | 22.27 | 0.306 |
| 47 | 1F15_C | 191 | α/β | 25.92 | 0.235 | 33.62 | 0.147 | 29.49 | 0.212 |
| 48 | 1F1E_A | 151 | α/β | 7.19 | 0.590 | 11.48 | 0.337 | 10.01 | 0.463 |
| 49 | 1F2R_I | 100 | α/β | 7.27 | 0.604 | 12.48 | 0.322 | 8.75 | 0.579 |
| 50 | 1F3U_A | 118 | α/β | 20.89 | 0.166 | 21.06 | 0.133 | 21.41 | 0.131 |
| 51 | 1F3Y_A | 165 | α/β | 6.07 | 0.737 | 19.16 | 0.318 | 7.00 | 0.697 |
| 52 | 1F43_A | 61 | α | 4.39 | 0.581 | 7.25 | 0.573 | 4.15 | 0.571 |
| 53 | 1F93_A | 103 | α/β | 2.08 | 0.801 | 8.53 | 0.478 | 2.14 | 0.796 |
| 54 | 1F98_A | 125 | α/β | 7.72 | 0.722 | 10.50 | 0.449 | 9.20 | 0.715 |
| 55 | 1F9P_A | 81 | α/β | 7.61 | 0.600 | 8.75 | 0.460 | 11.55 | 0.577 |
| 56 | 1FAQ_A | 52 | β | 2.73 | 0.601 | 9.21 | 0.320 | 2.98 | 0.561 |
| 57 | 1FC3_A | 116 | α | 2.76 | 0.752 | 9.32 | 0.373 | 3.01 | 0.743 |
| 58 | 1FCA_A | 55 | α/β | 1.43 | 0.783 | 3.31 | 0.533 | 1.55 | 0.755 |
| 59 | 1FEX_A | 59 | α | 2.45 | 0.620 | 8.57 | 0.306 | 2.55 | 0.608 |
| 60 | 1FHT_A | 116 | α/β | 8.53 | 0.689 | 17.04 | 0.385 | 7.93 | 0.675 |
| 61 | 1FJG_F | 101 | α/β | 3.38 | 0.809 | 11.20 | 0.389 | 3.38 | 0.809 |
| 62 | 1FJG_H | 138 | α/β | 2.67 | 0.811 | 13.96 | 0.328 | 3.19 | 0.794 |
| 63 | 1FMB_A | 104 | α/β | 5.16 | 0.721 | 13.10 | 0.276 | 5.42 | 0.696 |
| 64 | 1FR0_A | 125 | α | 4.17 | 0.671 | 10.00 | 0.412 | 4.63 | 0.646 |
| 65 | 1FRD_A | 98 | α/β | 2.48 | 0.748 | 10.00 | 0.287 | 2.66 | 0.734 |
| 66 | 1FSP_A | 124 | α/β | 3.40 | 0.819 | 3.73 | 0.776 | 3.32 | 0.811 |
| 67 | 1FW9_A | 164 | α/β | 3.32 | 0.758 | 14.40 | 0.394 | 6.17 | 0.726 |
| 68 | 1G2R_A | 94 | α/β | 2.01 | 0.810 | 12.99 | 0.294 | 2.04 | 0.799 |
| 69 | 1GGS_A | 81 | α/β | 6.25 | 0.655 | 8.91 | 0.635 | 10.39 | 0.637 |
| 70 | 1GME_A | 150 | α/β | 9.98 | 0.559 | 16.04 | 0.278 | 10.57 | 0.530 |
| 71 | 1GPQ_B | 128 | α/β | 5.19 | 0.663 | 12.93 | 0.361 | 4.85 | 0.646 |
| 72 | 1GQA_A | 130 | α/β | 2.60 | 0.799 | 10.90 | 0.389 | 2.76 | 0.777 |
| 73 | 1GVN_A | 87 | α | 17.00 | 0.235 | 18.65 | 0.277 | 16.79 | 0.221 |
| 74 | 1GVP_A | 87 | α/β | 5.36 | 0.689 | 9.99 | 0.319 | 5.46 | 0.684 |
| 75 | 1GXD_C | 192 | α/β | 13.02 | 0.530 | 13.87 | 0.341 | 13.60 | 0.497 |
| 76 | 1H8E_H | 89 | α/β | 1.66 | 0.840 | 12.65 | 0.230 | 1.82 | 0.822 |
| 77 | 1H9E_A | 56 | α | 4.11 | 0.519 | 6.91 | 0.373 | 4.53 | 0.493 |
| 78 | 1H9F_A | 57 | α | 4.30 | 0.516 | 10.95 | 0.395 | 4.56 | 0.502 |
| 79 | 1HBG_A | 147 | α | 2.09 | 0.855 | 12.52 | 0.320 | 2.06 | 0.853 |
| 80 | 1HBX_E | 89 | α/β | 16.43 | 0.509 | 12.79 | 0.453 | 16.32 | 0.469 |
| 81 | 1HCD_A | 118 | α/β | 3.20 | 0.718 | 10.50 | 0.283 | 3.25 | 0.712 |
| 82 | 1HH8_A | 192 | α/β | 12.60 | 0.695 | 16.25 | 0.577 | 19.26 | 0.656 |
| 83 | 1HHV_A | 74 | α/β | 5.72 | 0.590 | 7.26 | 0.452 | 13.33 | 0.565 |
| 84 | 1HKQ_A | 125 | α/β | 8.85 | 0.685 | 16.71 | 0.310 | 6.98 | 0.670 |
| 85 | 1HKX_E | 143 | α/β | 4.46 | 0.696 | 14.20 | 0.276 | 5.43 | 0.682 |
| 86 | 1HL6_D | 143 | α/β | 4.55 | 0.660 | 19.08 | 0.297 | 4.35 | 0.651 |

| NO. | PDB | Length | Type | GDDfold | | GDDfold_global | | GDDfold_local | |
|---|---|---|---|---|---|---|---|---|---|
| | | | | RMSD | TMscore | RMSD | TMscore | RMSD | TMscore |
| 87 | 1I35_A | 95 | α/β | 3.86 | 0.582 | 10.98 | 0.346 | 4.62 | 0.566 |
| 88 | 1I85_A | 110 | β | 2.22 | 0.789 | 10.92 | 0.409 | 2.41 | 0.768 |
| 89 | 1ID2_A | 106 | α/β | 5.60 | 0.738 | 15.38 | 0.307 | 5.22 | 0.724 |
| 90 | 1IM3_D | 95 | α/β | 12.28 | 0.351 | 19.61 | 0.234 | 14.08 | 0.260 |
| 91 | 1IOO_A | 196 | α/β | 4.02 | 0.740 | 17.11 | 0.312 | 4.34 | 0.712 |
| 92 | 1IRS_A | 112 | α/β | 2.62 | 0.811 | 12.12 | 0.287 | 2.62 | 0.811 |
| 93 | 1IS7_K | 85 | α/β | 5.45 | 0.622 | 12.83 | 0.286 | 6.68 | 0.582 |
| 94 | 1IUJ_B | 103 | α/β | 1.89 | 0.837 | 10.56 | 0.414 | 2.21 | 0.834 |
| 95 | 1IUY_A | 92 | α/β | 8.78 | 0.665 | 14.38 | 0.355 | 8.78 | 0.665 |
| 96 | 1J8I_A | 93 | α/β | 9.47 | 0.537 | 12.79 | 0.408 | 20.64 | 0.521 |
| 97 | 1J9I_A | 68 | α/β | 5.41 | 0.556 | 13.07 | 0.320 | 5.72 | 0.545 |
| 98 | 1JEI_A | 53 | α | 6.35 | 0.590 | 5.59 | 0.414 | 4.84 | 0.588 |
| 99 | 1JIW_I | 105 | α/β | 3.03 | 0.732 | 12.36 | 0.335 | 3.67 | 0.705 |
| 100 | 1JJ2_S | 119 | α/β | 4.53 | 0.698 | 9.06 | 0.411 | 5.63 | 0.656 |
| 101 | 1JLI_A | 112 | α | 12.71 | 0.355 | 15.81 | 0.237 | 12.77 | 0.326 |
| 102 | 1JMT_A | 98 | α/β | 4.64 | 0.731 | 10.96 | 0.327 | 4.43 | 0.703 |
| 103 | 1JO0_A | 97 | α/β | 2.05 | 0.826 | 6.62 | 0.556 | 2.51 | 0.816 |
| 104 | 1JOP_A | 140 | α/β | 3.13 | 0.741 | 15.13 | 0.294 | 3.43 | 0.728 |
| 105 | 1JPY_Y | 120 | α/β | 7.65 | 0.578 | 18.75 | 0.239 | 14.72 | 0.524 |
| 106 | 1JR5_A | 90 | α | 7.75 | 0.549 | 11.57 | 0.338 | 7.83 | 0.516 |
| 107 | 1K1Z_A | 78 | β | 3.97 | 0.589 | 13.26 | 0.336 | 7.11 | 0.533 |
| 108 | 1K3S_A | 109 | α/β | 3.23 | 0.689 | 10.06 | 0.393 | 3.34 | 0.675 |
| 109 | 1K5D_B | 146 | α/β | 9.67 | 0.753 | 18.38 | 0.283 | 11.30 | 0.739 |
| 110 | 1K73_1 | 73 | α/β | 5.56 | 0.539 | 7.65 | 0.320 | 5.54 | 0.523 |
| 111 | 1KA8_A | 100 | α/β | 6.31 | 0.570 | 13.69 | 0.287 | 6.52 | 0.551 |
| 112 | 1KN6_A | 73 | α/β | 3.33 | 0.615 | 9.39 | 0.342 | 3.43 | 0.592 |
| 113 | 1KOH_D | 172 | α/β | 6.63 | 0.751 | 15.28 | 0.344 | 7.55 | 0.740 |
| 114 | 1KP6_A | 79 | α/β | 13.25 | 0.285 | 13.80 | 0.250 | 15.02 | 0.218 |
| 115 | 1KPT_A | 105 | α/β | 3.30 | 0.689 | 12.52 | 0.301 | 3.62 | 0.660 |
| 116 | 1KQ6_A | 140 | α/β | 7.43 | 0.706 | 16.04 | 0.330 | 13.31 | 0.661 |
| 117 | 1KSX_A | 144 | α/β | 2.52 | 0.800 | 13.35 | 0.314 | 3.26 | 0.760 |
| 118 | 1KX5_D | 107 | α | 5.34 | 0.569 | 10.06 | 0.408 | 8.59 | 0.513 |
| 119 | 1L1D_B | 147 | α/β | 2.59 | 0.825 | 16.25 | 0.261 | 2.67 | 0.817 |
| 120 | 1L3G_A | 123 | α/β | 6.26 | 0.561 | 17.96 | 0.364 | 8.16 | 0.531 |
| 121 | 1L6H_A | 69 | α | 3.66 | 0.511 | 7.40 | 0.377 | 3.99 | 0.487 |
| 122 | 1LDD_A | 71 | α/β | 1.50 | 0.830 | 11.90 | 0.324 | 1.67 | 0.813 |
| 123 | 1LE2_A | 144 | α | 54.28 | 0.240 | 56.10 | 0.223 | 51.86 | 0.225 |
| 124 | 1LFU_P | 82 | α | 11.52 | 0.554 | 13.56 | 0.347 | 11.41 | 0.540 |
| 125 | 1LNW_C | 139 | α/β | 6.87 | 0.712 | 13.00 | 0.470 | 7.00 | 0.698 |
| 126 | 1LQM_H | 84 | α/β | 5.40 | 0.546 | 11.56 | 0.205 | 9.12 | 0.408 |
| 127 | 1LR1_B | 57 | α | 5.55 | 0.364 | 4.90 | 0.429 | 5.71 | 0.352 |
| 128 | 1LZW_B | 146 | α/β | 2.38 | 0.843 | 14.95 | 0.316 | 2.71 | 0.823 |
| 129 | 1MAI_A | 119 | α/β | 2.27 | 0.821 | 12.50 | 0.295 | 3.29 | 0.789 |
| 130 | 1MFQ_C | 108 | α | 9.60 | 0.588 | 13.24 | 0.468 | 9.83 | 0.583 |

| NO. | PDB | Length | Type | GDDfold | | GDDfold_global | | GDDfold_local | |
|---|---|---|---|---|---|---|---|---|---|
| | | | | RMSD | TMscore | RMSD | TMscore | RMSD | TMscore |
| 131 | 1MWQ_A | 99 | α/β | 2.55 | 0.768 | 8.00 | 0.332 | 3.12 | 0.752 |
| 132 | 1N12_A | 138 | α/β | 3.44 | 0.738 | 15.62 | 0.222 | 3.49 | 0.730 |
| 133 | 1N3G_A | 113 | α/β | 7.16 | 0.727 | 10.00 | 0.446 | 7.38 | 0.717 |
| 134 | 1NF6_F | 171 | α/β | 3.03 | 0.774 | 16.35 | 0.335 | 7.38 | 0.740 |
| 135 | 1NGL_A | 179 | α/β | 7.01 | 0.662 | 13.74 | 0.427 | 9.26 | 0.645 |
| 136 | 1NKZ_A | 53 | α | 3.48 | 0.557 | 7.71 | 0.494 | 5.78 | 0.399 |
| 137 | 1NLQ_A | 105 | α/β | 2.34 | 0.803 | 13.69 | 0.268 | 2.77 | 0.779 |
| 138 | 1NOE_A | 86 | α/β | 4.44 | 0.605 | 12.02 | 0.266 | 4.44 | 0.605 |
| 139 | 1NPB_A | 140 | α/β | 7.82 | 0.678 | 14.36 | 0.359 | 8.75 | 0.653 |
| 140 | 1NQZ_A | 171 | α/β | 6.20 | 0.784 | 16.83 | 0.367 | 7.19 | 0.765 |
| 141 | 1NR3_A | 122 | α/β | 17.46 | 0.319 | 14.89 | 0.241 | 15.75 | 0.305 |
| 142 | 1NTV_A | 152 | α/β | 4.01 | 0.758 | 17.67 | 0.280 | 4.46 | 0.751 |
| 143 | 1NZE_A | 112 | α | 3.58 | 0.839 | 9.95 | 0.383 | 3.78 | 0.834 |
| 144 | 1O0G_A | 124 | α/β | 4.78 | 0.646 | 15.28 | 0.287 | 6.26 | 0.604 |
| 145 | 1OA8_D | 133 | α/β | 13.11 | 0.369 | 21.82 | 0.196 | 14.59 | 0.308 |
| 146 | 1OFT_A | 119 | α/β | 2.52 | 0.791 | 7.12 | 0.464 | 2.59 | 0.785 |
| 147 | 1OJG_A | 136 | α/β | 3.93 | 0.635 | 6.90 | 0.513 | 4.03 | 0.630 |
| 148 | 1OOF_A | 124 | α/β | 2.11 | 0.830 | 7.37 | 0.420 | 2.33 | 0.809 |
| 149 | 1ORY_A | 119 | α | 3.58 | 0.756 | 9.53 | 0.329 | 3.91 | 0.740 |
| 150 | 1OX7_A | 158 | α/β | 3.08 | 0.795 | 9.09 | 0.440 | 3.88 | 0.782 |
| 151 | 1OZ9_A | 141 | α/β | 2.62 | 0.792 | 8.41 | 0.555 | 4.04 | 0.745 |
| 152 | 1PD6_A | 94 | β | 5.14 | 0.629 | 8.51 | 0.418 | 5.09 | 0.608 |
| 153 | 1PFS_A | 78 | β | 4.97 | 0.573 | 11.13 | 0.232 | 5.70 | 0.554 |
| 154 | 1PGV_A | 167 | α/β | 3.23 | 0.773 | 4.69 | 0.674 | 3.56 | 0.750 |
| 155 | 1PIH_A | 73 | α/β | 4.53 | 0.620 | 10.09 | 0.272 | 5.72 | 0.586 |
| 156 | 1PMS_A | 135 | α/β | 5.83 | 0.632 | 11.80 | 0.325 | 8.48 | 0.619 |
| 157 | 1PSR_A | 100 | α/β | 4.80 | 0.677 | 7.29 | 0.374 | 4.69 | 0.669 |
| 158 | 1PXW_A | 128 | α/β | 5.57 | 0.717 | 17.89 | 0.361 | 6.91 | 0.706 |
| 159 | 1PZW_A | 80 | α/β | 2.52 | 0.731 | 13.78 | 0.294 | 2.76 | 0.725 |
| 160 | 1QFT_A | 175 | α/β | 3.94 | 0.773 | 12.71 | 0.402 | 4.39 | 0.760 |
| 161 | 1QFW_B | 110 | β | 7.77 | 0.576 | 12.58 | 0.287 | 10.81 | 0.512 |
| 162 | 1QMA_A | 123 | α/β | 1.74 | 0.877 | 11.72 | 0.410 | 1.93 | 0.868 |
| 163 | 1QZG_A | 170 | α/β | 5.15 | 0.700 | 12.85 | 0.283 | 4.14 | 0.698 |
| 164 | 1R5T_A | 141 | α/β | 3.10 | 0.810 | 13.65 | 0.286 | 3.28 | 0.808 |
| 165 | 1R6R_A | 80 | α | 8.42 | 0.460 | 9.63 | 0.355 | 7.09 | 0.436 |
| 166 | 1RHX_A | 87 | α/β | 4.28 | 0.615 | 9.27 | 0.338 | 4.37 | 0.601 |
| 167 | 1ROW_A | 107 | α/β | 2.03 | 0.819 | 12.33 | 0.298 | 2.15 | 0.799 |
| 168 | 1RTU_A | 114 | α/β | 3.48 | 0.696 | 16.88 | 0.241 | 3.46 | 0.692 |
| 169 | 1RZ3_A | 184 | α/β | 3.52 | 0.817 | 13.18 | 0.302 | 4.81 | 0.769 |
| 170 | 1S2D_A | 165 | α/β | 5.30 | 0.763 | 8.33 | 0.456 | 5.38 | 0.735 |
| 171 | 1S3J_A | 143 | α/β | 4.83 | 0.638 | 13.07 | 0.479 | 5.89 | 0.614 |
| 172 | 1S56_B | 135 | α/β | 6.68 | 0.754 | 17.04 | 0.271 | 5.99 | 0.736 |
| 173 | 1S7O_C | 108 | α | 8.07 | 0.540 | 7.04 | 0.512 | 8.47 | 0.530 |
| 174 | 1S7Z_A | 106 | α | 4.83 | 0.629 | 10.33 | 0.400 | 6.65 | 0.597 |

| NO. | PDB | Length | Type | GDDfold | | GDDfold_global | | GDDfold_local | |
|---|---|---|---|---|---|---|---|---|---|
| | | | | RMSD | TMscore | RMSD | TMscore | RMSD | TMscore |
| 175 | 1SAU_A | 114 | α/β | 3.01 | 0.709 | 12.94 | 0.298 | 3.17 | 0.688 |
| 176 | 1SMP_I | 100 | α/β | 2.53 | 0.764 | 12.72 | 0.386 | 2.59 | 0.756 |
| 177 | 1SPP_B | 112 | α/β | 2.70 | 0.762 | 12.85 | 0.282 | 3.84 | 0.719 |
| 178 | 1STM_A | 141 | α/β | 5.35 | 0.594 | 17.12 | 0.239 | 7.12 | 0.552 |
| 179 | 1SVJ_A | 136 | α/β | 2.58 | 0.787 | 11.43 | 0.367 | 3.19 | 0.754 |
| 180 | 1TEO_A | 173 | α/β | 4.96 | 0.718 | 14.89 | 0.391 | 5.05 | 0.718 |
| 181 | 1TJF_B | 186 | α/β | 3.98 | 0.837 | 10.57 | 0.483 | 3.98 | 0.837 |
| 182 | 1TLJ_A | 189 | α/β | 3.45 | 0.778 | 14.33 | 0.362 | 3.90 | 0.741 |
| 183 | 1TUL_A | 102 | α/β | 3.98 | 0.666 | 13.20 | 0.243 | 3.98 | 0.666 |
| 184 | 1TWU_A | 137 | α/β | 4.13 | 0.716 | 12.17 | 0.385 | 3.98 | 0.708 |
| 185 | 1TYG_B | 65 | α/β | 2.06 | 0.755 | 7.04 | 0.373 | 2.12 | 0.749 |
| 186 | 1TZ0_A | 108 | α/β | 3.18 | 0.717 | 14.81 | 0.269 | 4.22 | 0.684 |
| 187 | 1U84_A | 81 | α | 1.89 | 0.809 | 4.69 | 0.580 | 1.89 | 0.809 |
| 188 | 1UFB_A | 127 | α | 2.38 | 0.817 | 12.37 | 0.371 | 2.74 | 0.792 |
| 189 | 1UG4_A | 60 | β | 5.74 | 0.623 | 9.65 | 0.337 | 4.07 | 0.610 |
| 190 | 1UNG_D | 149 | α | 4.17 | 0.705 | 13.28 | 0.310 | 4.64 | 0.644 |
| 191 | 1USL_C | 158 | α/β | 3.93 | 0.851 | 9.78 | 0.551 | 4.06 | 0.845 |
| 192 | 1V74_A | 107 | α/β | 6.42 | 0.639 | 11.19 | 0.436 | 5.29 | 0.630 |
| 193 | 1VCC_A | 77 | α/β | 5.59 | 0.509 | 10.88 | 0.251 | 7.75 | 0.446 |
| 194 | 1VCY_A | 193 | α/β | 4.29 | 0.735 | 19.74 | 0.284 | 5.90 | 0.713 |
| 195 | 1VD0_A | 109 | α/β | 12.54 | 0.651 | 16.64 | 0.297 | 12.14 | 0.639 |
| 196 | 1VHG_A | 185 | α/β | 12.70 | 0.665 | 13.99 | 0.406 | 10.95 | 0.652 |
| 197 | 1VKE_E | 119 | α | 10.91 | 0.643 | 14.08 | 0.481 | 11.08 | 0.591 |
| 198 | 1VYI_A | 111 | α/β | 21.15 | 0.232 | 22.25 | 0.277 | 21.48 | 0.209 |
| 199 | 1VYX_A | 60 | α/β | 5.78 | 0.457 | 9.46 | 0.273 | 6.13 | 0.440 |
| 200 | 1W1W_E | 70 | α/β | 2.25 | 0.782 | 4.39 | 0.479 | 2.44 | 0.753 |
| 201 | 1WJ8_A | 117 | α | 1.79 | 0.860 | 8.55 | 0.439 | 1.82 | 0.852 |
| 202 | 1WLQ_C | 185 | α/β | 6.92 | 0.640 | 14.14 | 0.342 | 7.56 | 0.594 |
| 203 | 1WMH_B | 82 | α/β | 1.70 | 0.832 | 10.82 | 0.343 | 1.84 | 0.814 |
| 204 | 1XJA_C | 169 | α/β | 4.15 | 0.748 | 19.71 | 0.358 | 7.02 | 0.718 |
| 205 | 1Y14_A | 133 | α | 9.56 | 0.525 | 12.87 | 0.295 | 9.45 | 0.497 |
| 206 | 1Y1X_A | 182 | α/β | 5.23 | 0.653 | 11.65 | 0.420 | 6.37 | 0.625 |
| 207 | 1YG2_A | 169 | α/β | 11.50 | 0.483 | 21.57 | 0.328 | 18.14 | 0.432 |
| 208 | 1Z8R_A | 150 | α/β | 12.46 | 0.471 | 19.62 | 0.185 | 13.10 | 0.433 |
| 209 | 2A5Y_A | 173 | α | 3.95 | 0.744 | 17.46 | 0.415 | 4.92 | 0.714 |
| 210 | 2A9U_B | 127 | α | 11.29 | 0.658 | 10.90 | 0.384 | 12.78 | 0.642 |
| 211 | 2ACY_A | 98 | α/β | 3.16 | 0.793 | 13.62 | 0.316 | 2.85 | 0.778 |
| 212 | 2AEN_A | 164 | α/β | 6.27 | 0.621 | 18.28 | 0.228 | 12.31 | 0.509 |
| 213 | 2APN_A | 114 | α/β | 7.98 | 0.651 | 9.14 | 0.466 | 8.39 | 0.621 |
| 214 | 2AQ0_A | 84 | α | 7.10 | 0.637 | 7.94 | 0.514 | 7.08 | 0.611 |
| 215 | 2AQS_A | 160 | α/β | 4.11 | 0.766 | 15.23 | 0.297 | 4.92 | 0.743 |
| 216 | 2BSE_A | 107 | α/β | 4.27 | 0.601 | 17.07 | 0.236 | 6.61 | 0.563 |
| 217 | 2BWJ_A | 196 | α/β | 4.06 | 0.780 | 13.82 | 0.405 | 5.98 | 0.754 |
| 218 | 2BYK_D | 92 | α | 4.42 | 0.586 | 10.53 | 0.344 | 5.28 | 0.562 |

| NO. | PDB | Length | Type | GDDfold | | GDDfold_global | | GDDfold_local | |
|---|---|---|---|---|---|---|---|---|---|
| | | | | RMSD | TMscore | RMSD | TMscore | RMSD | TMscore |
| 219 | 2C2F_A | 178 | α/β | 3.76 | 0.807 | 11.03 | 0.476 | 4.76 | 0.789 |
| 220 | 2C4W_A | 168 | α/β | 2.84 | 0.831 | 13.62 | 0.303 | 4.60 | 0.816 |
| 221 | 2CDP_A | 138 | α/β | 3.07 | 0.784 | 15.14 | 0.279 | 3.51 | 0.751 |
| 222 | 2CMX_A | 70 | α/β | 2.06 | 0.740 | 11.53 | 0.346 | 2.21 | 0.720 |
| 223 | 2CO3_B | 135 | α/β | 6.35 | 0.670 | 13.69 | 0.248 | 6.03 | 0.628 |
| 224 | 2CWP_A | 109 | α/β | 5.73 | 0.733 | 14.97 | 0.272 | 5.88 | 0.725 |
| 225 | 2CZV_D | 119 | α/β | 2.37 | 0.816 | 13.01 | 0.293 | 2.37 | 0.816 |
| 226 | 2D0P_B | 110 | α/β | 2.94 | 0.794 | 7.52 | 0.499 | 2.82 | 0.774 |
| 227 | 2EWC_B | 122 | α/β | 3.42 | 0.765 | 9.65 | 0.312 | 3.93 | 0.758 |
| 228 | 2F22_B | 143 | α/β | 3.14 | 0.789 | 13.81 | 0.294 | 3.26 | 0.759 |
| 229 | 2FA5_B | 142 | α/β | 7.84 | 0.723 | 11.64 | 0.539 | 8.37 | 0.683 |
| 230 | 2FKB_C | 167 | α/β | 2.96 | 0.804 | 11.20 | 0.402 | 4.40 | 0.780 |
| 231 | 2GBJ_B | 84 | α/β | 2.58 | 0.756 | 7.16 | 0.449 | 3.12 | 0.739 |
| 232 | 2GJ3_A | 119 | α/β | 2.70 | 0.805 | 8.41 | 0.434 | 2.84 | 0.790 |
| 233 | 2GKC_A | 155 | α/β | 4.97 | 0.629 | 13.32 | 0.307 | 4.86 | 0.619 |
| 234 | 2H30_A | 151 | α/β | 4.73 | 0.702 | 13.82 | 0.318 | 5.33 | 0.681 |
| 235 | 2H8E_A | 120 | α/β | 2.80 | 0.770 | 15.82 | 0.296 | 2.78 | 0.766 |
| 236 | 2HI3_A | 73 | α | 5.70 | 0.616 | 8.70 | 0.473 | 5.66 | 0.578 |
| 237 | 2HQ7_B | 142 | α/β | 2.98 | 0.797 | 12.08 | 0.386 | 3.95 | 0.755 |
| 238 | 2HYB_A | 130 | α/β | 2.15 | 0.830 | 14.93 | 0.244 | 3.47 | 0.777 |
| 239 | 2ICT_A | 94 | α | 7.48 | 0.729 | 10.50 | 0.525 | 5.70 | 0.713 |
| 240 | 2J4H_B | 172 | α/β | 7.55 | 0.727 | 17.34 | 0.275 | 9.71 | 0.699 |
| 241 | 2J6Z_A | 86 | α | 3.72 | 0.640 | 6.31 | 0.479 | 4.47 | 0.593 |
| 242 | 2JLP_D | 169 | α/β | 5.30 | 0.710 | 16.19 | 0.231 | 13.34 | 0.303 |
| 243 | 2JP3_A | 67 | α | 10.11 | 0.307 | 10.20 | 0.305 | 9.21 | 0.296 |
| 244 | 2K9X_A | 102 | α/β | 4.53 | 0.677 | 10.02 | 0.242 | 4.68 | 0.672 |
| 245 | 2KBW_A | 160 | α | 3.45 | 0.742 | 10.91 | 0.388 | 4.34 | 0.701 |
| 246 | 2L5P_A | 175 | α/β | 4.23 | 0.758 | 18.73 | 0.404 | 5.59 | 0.717 |
| 247 | 2L74_A | 125 | α/β | 6.95 | 0.702 | 10.14 | 0.448 | 9.22 | 0.669 |
| 248 | 2LKP_A | 119 | α/β | 9.48 | 0.602 | 13.69 | 0.343 | 10.03 | 0.601 |
| 249 | 2LRB_A | 165 | α/β | 3.95 | 0.720 | 15.75 | 0.282 | 3.95 | 0.708 |
| 250 | 2LWP_A | 97 | α/β | 4.90 | 0.550 | 8.58 | 0.315 | 7.40 | 0.525 |
| 251 | 2NAZ_A | 109 | α/β | 4.62 | 0.662 | 7.48 | 0.363 | 5.78 | 0.641 |
| 252 | 2NCM_A | 99 | β | 1.77 | 0.834 | 9.49 | 0.352 | 1.82 | 0.831 |
| 253 | 2NDP_A | 99 | α/β | 8.44 | 0.508 | 9.31 | 0.335 | 9.35 | 0.461 |
| 254 | 2NS9_B | 152 | α/β | 4.02 | 0.814 | 7.59 | 0.421 | 3.61 | 0.809 |
| 255 | 2O70_F | 168 | α | 2.74 | 0.795 | 12.11 | 0.296 | 2.80 | 0.781 |
| 256 | 2ODM_B | 83 | α | 2.93 | 0.709 | 6.73 | 0.426 | 3.28 | 0.693 |
| 257 | 2P7L_A | 125 | α/β | 4.17 | 0.749 | 8.98 | 0.403 | 4.66 | 0.730 |
| 258 | 2PI2_F | 119 | α/β | 3.26 | 0.761 | 15.70 | 0.391 | 4.82 | 0.708 |
| 259 | 2PYB_A | 151 | α | 3.39 | 0.817 | 12.32 | 0.344 | 3.42 | 0.816 |
| 260 | 2Q2H_A | 118 | α/β | 9.37 | 0.637 | 15.52 | 0.208 | 9.89 | 0.630 |
| 261 | 2QVG_A | 129 | α/β | 2.23 | 0.819 | 6.37 | 0.501 | 2.33 | 0.809 |
| 262 | 2QZJ_A | 121 | α/β | 1.54 | 0.889 | 2.54 | 0.783 | 1.58 | 0.884 |

| NO. | PDB | Length | Type | GDDfold | | GDDfold_global | | GDDfold_local | |
|---|---|---|---|---|---|---|---|---|---|
| | | | | RMSD | TMscore | RMSD | TMscore | RMSD | TMscore |
| 263 | 2RD5_D | 126 | α/β | 10.74 | 0.646 | 15.11 | 0.270 | 11.46 | 0.570 |
| 264 | 2RLD_C | 116 | α | 2.39 | 0.824 | 8.37 | 0.404 | 2.78 | 0.809 |
| 265 | 2UUX_A | 55 | α/β | 3.60 | 0.649 | 11.81 | 0.257 | 3.65 | 0.557 |
| 266 | 2V85_A | 74 | α/β | 9.08 | 0.379 | 10.30 | 0.319 | 10.12 | 0.287 |
| 267 | 2VUL_A | 193 | α/β | 2.54 | 0.830 | 11.76 | 0.417 | 2.54 | 0.830 |
| 268 | 2WCW_B | 122 | α/β | 3.25 | 0.796 | 12.10 | 0.289 | 3.30 | 0.781 |
| 269 | 2WGP_A | 168 | α/β | 2.85 | 0.809 | 8.66 | 0.569 | 4.62 | 0.780 |
| 270 | 2XGY_A | 129 | α | 5.69 | 0.684 | 10.85 | 0.359 | 7.97 | 0.617 |
| 271 | 2Z3B_A | 180 | α/β | 2.44 | 0.842 | 8.88 | 0.516 | 2.59 | 0.832 |
| 272 | 2ZMZ_B | 79 | α/β | 4.58 | 0.697 | 8.90 | 0.381 | 5.05 | 0.665 |
| 273 | 3ALU_A | 157 | α/β | 3.85 | 0.733 | 15.46 | 0.260 | 5.54 | 0.715 |
| 274 | 3BDB_A | 126 | α/β | 6.60 | 0.733 | 12.93 | 0.299 | 5.46 | 0.708 |
| 275 | 3CAE_A | 132 | α/β | 27.29 | 0.504 | 21.35 | 0.218 | 26.52 | 0.480 |
| 276 | 3CG4_A | 126 | α/β | 2.69 | 0.822 | 3.85 | 0.654 | 2.70 | 0.818 |
| 277 | 3CX5_F | 74 | α | 2.92 | 0.637 | 5.18 | 0.448 | 3.37 | 0.608 |
| 278 | 3CX5_G | 126 | α | 7.28 | 0.640 | 9.42 | 0.357 | 7.67 | 0.627 |
| 279 | 3E6M_E | 147 | α/β | 7.95 | 0.709 | 15.14 | 0.523 | 7.89 | 0.692 |
| 280 | 3E9T_D | 102 | β | 1.77 | 0.849 | 12.29 | 0.248 | 1.93 | 0.828 |
| 281 | 3EOD_A | 115 | α/β | 2.16 | 0.834 | 6.58 | 0.652 | 3.03 | 0.808 |
| 282 | 3F8L_A | 162 | α/β | 3.03 | 0.814 | 14.63 | 0.377 | 3.03 | 0.814 |
| 283 | 3G20_B | 119 | α/β | 7.46 | 0.642 | 12.46 | 0.384 | 7.14 | 0.626 |
| 284 | 3GMX_A | 153 | α/β | 14.33 | 0.410 | 14.85 | 0.293 | 13.45 | 0.393 |
| 285 | 3H05_B | 163 | α/β | 3.68 | 0.801 | 11.14 | 0.333 | 4.08 | 0.799 |
| 286 | 3I9V_7 | 127 | α/β | 5.20 | 0.626 | 15.79 | 0.258 | 6.33 | 0.563 |
| 287 | 3IAM_2 | 179 | α/β | 4.23 | 0.715 | 14.27 | 0.326 | 5.93 | 0.592 |
| 288 | 3LQV_B | 115 | α/β | 5.20 | 0.675 | 11.72 | 0.372 | 9.73 | 0.637 |
| 289 | 3M1N_B | 168 | α/β | 9.63 | 0.510 | 13.50 | 0.355 | 19.21 | 0.475 |
| 290 | 3MQK_C | 75 | β | 1.74 | 0.800 | 7.31 | 0.407 | 1.82 | 0.790 |
| 291 | 3N1G_C | 104 | α/β | 2.18 | 0.818 | 11.59 | 0.399 | 2.93 | 0.791 |
| 292 | 3N9U_C | 96 | α/β | 5.84 | 0.737 | 6.89 | 0.367 | 6.25 | 0.724 |
| 293 | 3O61_A | 187 | α/β | 13.05 | 0.690 | 16.50 | 0.298 | 14.21 | 0.670 |
| 294 | 3P8B_A | 60 | α/β | 2.17 | 0.663 | 8.36 | 0.342 | 2.50 | 0.608 |
| 295 | 3PD2_A | 147 | α/β | 3.23 | 0.776 | 18.05 | 0.274 | 3.67 | 0.730 |
| 296 | 3QU3_A | 122 | α/β | 4.29 | 0.656 | 13.69 | 0.245 | 4.34 | 0.650 |
| 297 | 3SDL_B | 97 | α | 31.18 | 0.237 | 26.45 | 0.185 | 20.10 | 0.232 |
| 298 | 3UE6_E | 138 | α/β | 5.65 | 0.729 | 13.19 | 0.385 | 5.45 | 0.721 |
| 299 | 3V1O_A | 165 | α/β | 4.25 | 0.762 | 15.77 | 0.283 | 6.42 | 0.726 |
| 300 | 3W1Z_D | 110 | α/β | 3.62 | 0.739 | 11.65 | 0.313 | 4.57 | 0.717 |
| 301 | 3X0G_A | 93 | α | 3.32 | 0.665 | 15.79 | 0.320 | 3.51 | 0.640 |
| 302 | 3X15_A | 87 | α | 15.06 | 0.419 | 12.63 | 0.312 | 20.25 | 0.376 |
| 303 | 4AIH_A | 139 | α/β | 4.22 | 0.745 | 8.34 | 0.414 | 6.55 | 0.713 |
| 304 | 4ASW_C | 81 | α/β | 1.95 | 0.796 | 7.15 | 0.500 | 2.10 | 0.781 |
| 305 | 4B0M_A | 131 | α/β | 4.52 | 0.718 | 11.98 | 0.360 | 11.97 | 0.627 |
| 306 | 4CXT_A | 132 | α/β | 6.22 | 0.728 | 15.08 | 0.363 | 6.99 | 0.707 |

| NO. | PDB | Length | Type | GDDfold | | GDDfold_global | | GDDfold_local | |
|---|---|---|---|---|---|---|---|---|---|
| | | | | RMSD | TMscore | RMSD | TMscore | RMSD | TMscore |
| 307 | 4DYW_A | 129 | α/β | 1.84 | 0.865 | 13.96 | 0.394 | 2.12 | 0.845 |
| 308 | 4ESB_A | 103 | α/β | 5.57 | 0.745 | 8.33 | 0.358 | 5.46 | 0.724 |
| 309 | 4GDK_A | 88 | α/β | 3.23 | 0.770 | 8.51 | 0.299 | 2.67 | 0.750 |
| 310 | 4GF3_A | 123 | α/β | 7.13 | 0.696 | 12.82 | 0.360 | 7.16 | 0.679 |
| 311 | 4GQY_A | 147 | α/β | 8.77 | 0.747 | 12.33 | 0.276 | 7.32 | 0.728 |
| 312 | 4I60_A | 128 | α/β | 3.01 | 0.777 | 8.81 | 0.455 | 3.44 | 0.737 |
| 313 | 4IOS_A | 100 | α/β | 2.76 | 0.707 | 14.79 | 0.283 | 3.32 | 0.660 |
| 314 | 4J20_A | 88 | α/β | 2.68 | 0.746 | 10.22 | 0.343 | 2.72 | 0.725 |
| 315 | 4JGX_B | 128 | α/β | 3.76 | 0.718 | 16.18 | 0.286 | 4.22 | 0.697 |
| 316 | 4K1F_A | 198 | α/β | 3.65 | 0.781 | 10.91 | 0.432 | 4.33 | 0.771 |
| 317 | 4KA0_A | 143 | α/β | 2.25 | 0.852 | 14.30 | 0.338 | 2.27 | 0.840 |
| 318 | 4LE0_B | 133 | α/β | 1.88 | 0.874 | 4.11 | 0.723 | 1.90 | 0.869 |
| 319 | 4LMS_A | 80 | α/β | 8.25 | 0.462 | 10.50 | 0.287 | 13.53 | 0.308 |
| 320 | 4M75_F | 75 | α/β | 4.59 | 0.759 | 8.70 | 0.416 | 3.98 | 0.743 |
| 321 | 4MLF_D | 61 | β | 14.00 | 0.286 | 10.32 | 0.211 | 15.75 | 0.245 |
| 322 | 4MMG_A | 91 | α/β | 2.06 | 0.813 | 14.34 | 0.311 | 2.20 | 0.794 |
| 323 | 4NBI_A | 163 | α/β | 3.32 | 0.810 | 15.65 | 0.339 | 3.21 | 0.797 |
| 324 | 4OW1_A | 86 | α/β | 3.05 | 0.713 | 9.08 | 0.339 | 3.05 | 0.681 |
| 325 | 4Q2O_A | 92 | α/β | 5.04 | 0.768 | 13.18 | 0.336 | 5.68 | 0.746 |
| 326 | 4Q2Q_A | 90 | α/β | 2.80 | 0.786 | 7.21 | 0.503 | 6.89 | 0.765 |
| 327 | 4R67_0 | 199 | α/β | 2.63 | 0.851 | 7.75 | 0.591 | 2.79 | 0.840 |
| 328 | 4RUV_A | 106 | α/β | 1.87 | 0.832 | 7.90 | 0.542 | 1.95 | 0.822 |
| 329 | 4UIJ_A | 104 | α/β | 1.95 | 0.814 | 7.30 | 0.405 | 2.34 | 0.786 |
| 330 | 4V2O_A | 78 | α | 3.05 | 0.653 | 7.94 | 0.466 | 4.17 | 0.555 |
| 331 | 4Z6J_A | 133 | β | 3.04 | 0.738 | 9.62 | 0.385 | 3.43 | 0.723 |
| 332 | 4ZBY_A | 194 | α/β | 2.21 | 0.868 | 6.38 | 0.541 | 2.35 | 0.856 |
| 333 | 5CJ3_B | 126 | α/β | 3.42 | 0.789 | 13.53 | 0.325 | 3.39 | 0.767 |
| 334 | 5E4E_A | 111 | α/β | 8.80 | 0.384 | 18.66 | 0.203 | 9.22 | 0.328 |
| 335 | 5EKT_A | 196 | α/β | 3.36 | 0.834 | 9.95 | 0.409 | 3.47 | 0.827 |
| 336 | 5IAO_A | 171 | α/β | 3.75 | 0.773 | 9.24 | 0.472 | 4.15 | 0.742 |
| 337 | 5IZB_A | 89 | α/β | 5.75 | 0.567 | 18.41 | 0.403 | 11.97 | 0.516 |
| 338 | 5JTM_A | 155 | α/β | 10.91 | 0.703 | 15.70 | 0.300 | 10.79 | 0.693 |
| 339 | 5L38_A | 91 | α/β | 2.17 | 0.825 | 11.50 | 0.310 | 3.19 | 0.816 |
| 340 | 5L8R_D | 143 | α/β | 12.72 | 0.560 | 15.19 | 0.290 | 10.47 | 0.530 |
| 341 | 5O2V_A | 92 | α/β | 4.96 | 0.755 | 6.18 | 0.458 | 5.39 | 0.728 |
| 342 | 5O8G_A | 122 | α/β | 3.46 | 0.741 | 11.02 | 0.328 | 6.73 | 0.714 |
| 343 | 5T17_A | 85 | α/β | 3.13 | 0.655 | 4.60 | 0.495 | 3.18 | 0.635 |
| 344 | 5TMF_E | 95 | α/β | 7.64 | 0.545 | 11.50 | 0.316 | 9.52 | 0.505 |
| 345 | 5TUV_B | 104 | α/β | 13.95 | 0.377 | 15.05 | 0.345 | 11.44 | 0.302 |
| 346 | 5WSE_A | 114 | α/β | 3.66 | 0.792 | 8.89 | 0.578 | 3.97 | 0.787 |
| 347 | 6AQ3_B | 171 | α/β | 3.56 | 0.752 | 10.96 | 0.365 | 3.96 | 0.717 |

**Table S4**. Prediction results (TM-score) of GDDfold, GDDfold_relax, QUARK, RaptorX-DeepModeller, BAKER-ROSETTASERVER and MULTICOM_ CLUSTER for 24 free modeling FM targets of CASP13.

| No. | PDB | Length | GDDfold | GDDfold _relax | Quark | RaptorX-Deep Modeller | BAKER-ROSETTA SERVER | MULTI COM_ CLUSTER |
|---|---|---|---|---|---|---|---|---|
| 1 | T0950-D1 | 342 | 0.57 | 0.49 | 0.51 | 0.59 | 0.46 | 0.24 |
| 2 | T0953s1-D1 | 67 | 0.40 | 0.44 | 0.43 | 0.31 | 0.44 | 0.39 |
| 3 | T0953s2-D1 | 44 | 0.26 | 0.34 | 0.46 | 0.24 | 0.40 | 0.29 |
| 4 | T0953s2-D2 | 111 | 0.58 | 0.63 | 0.51 | 0.69 | 0.47 | 0.29 |
| 5 | T0953s2-D3 | 93 | 0.21 | 0.17 | 0.35 | 0.30 | 0.22 | 0.30 |
| 6 | T0955-D1 | 41 | 0.35 | 0.52 | 0.82 | 0.59 | \ | 0.79 |
| 7 | T0957s1-D1 | 108 | 0.53 | 0.56 | 0.46 | 0.37 | 0.42 | 0.31 |
| 8 | T0957s2-D1 | 155 | 0.68 | 0.76 | 0.63 | 0.65 | 0.54 | 0.51 |
| 9 | T0958-D1 | 77 | 0.73 | 0.73 | 0.55 | 0.67 | 0.62 | 0.56 |
| 10 | T0960-D2 | 84 | 0.46 | 0.52 | 0.44 | 0.49 | 0.54 | 0.39 |
| 11 | T0963-D2 | 82 | 0.48 | 0.53 | 0.46 | 0.54 | 0.36 | 0.64 |
| 12 | T0968s1-D1 | 118 | 0.71 | 0.71 | 0.57 | 0.62 | 0.74 | 0.43 |
| 13 | T0968s2-D1 | 115 | 0.67 | 0.75 | 0.65 | 0.67 | 0.78 | 0.42 |
| 14 | T0969-D1 | 354 | 0.78 | 0.81 | 0.64 | 0.66 | 0.49 | 0.46 |
| 15 | T0970-D1 | 85 | 0.69 | 0.71 | 0.58 | 0.54 | 0.57 | 0.33 |
| 16 | T0980s1-D1 | 104 | 0.56 | 0.64 | 0.54 | 0.48 | 0.41 | 0.27 |
| 17 | T0990-D1 | 76 | 0.56 | 0.63 | 0.65 | 0.40 | 0.37 | 0.37 |
| 18 | T0990-D2 | 231 | 0.48 | 0.49 | 0.37 | 0.35 | 0.26 | 0.27 |
| 19 | T0990-D3 | 213 | 0.36 | 0.36 | 0.27 | 0.34 | 0.24 | 0.24 |
| 20 | T1005-D1 | 326 | 0.70 | 0.69 | 0.71 | 0.72 | 0.72 | 0.69 |
| 21 | T1008-D1 | 77 | 0.52 | 0.69 | 0.66 | 0.31 | 0.65 | 0.56 |
| 22 | T1021s3-D1 | 166 | 0.73 | 0.70 | 0.64 | 0.67 | 0.50 | 0.50 |
| 23 | T1021s3-D2 | 97 | 0.47 | 0.53 | 0.45 | 0.59 | 0.22 | 0.27 |
| 24 | T1022s1-D1 | 156 | 0.54 | 0.58 | 0.66 | 0.60 | 0.42 | 0.49 |

**Table S5**. Prediction results (TM-score) of GDDfold, GDDfold_relax, QUARK, RaptorX, BAKER-ROSETTASERVER, MULTICOM_CLUSTER, and Yang-Server for 20 free modeling FM targets of CASP14.

| No. | PDB | Length | GDDfold | GDDfold _relax | Quark | RaptorX -Deep Modeller | BAKER-ROSETTA SERVER | MULTI COM_ CLUSTER |
|---|---|---|---|---|---|---|---|---|
| 1 | T1027-D1 | 99 | 0.38 | 0.37 | 0.47 | 0.41 | 0.34 | 0.37 |
| 2 | T1029-D1 | 125 | 0.47 | 0.45 | 0.47 | 0.45 | 0.49 | 0.46 |
| 3 | T1031-D1 | 95 | 0.23 | 0.24 | 0.72 | 0.35 | 0.24 | 0.35 |
| 4 | T1033-D1 | 100 | 0.41 | 0.31 | 0.36 | 0.33 | 0.44 | 0.41 |
| 5 | T1035-D1 | 102 | 0.47 | 0.60 | 0.83 | 0.43 | 0.26 | 0.77 |
| 6 | T1037-D1 | 404 | 0.54 | 0.68 | 0.79 | 0.53 | 0.32 | 0.73 |
| 7 | T1038-D1 | 114 | 0.36 | 0.36 | 0.36 | 0.39 | 0.30 | 0.35 |
| 8 | T1038-D2 | 76 | 0.69 | 0.66 | 0.60 | 0.54 | 0.62 | 0.57 |
| 9 | T1039-D1 | 161 | 0.29 | 0.32 | 0.42 | 0.42 | 0.48 | 0.23 |
| 10 | T1040-D1 | 130 | 0.26 | 0.25 | 0.43 | 0.30 | 0.25 | 0.26 |
| 11 | T1041-D1 | 242 | 0.71 | 0.71 | 0.76 | 0.70 | 0.41 | 0.65 |
| 12 | T1042-D1 | 276 | 0.20 | 0.24 | 0.72 | 0.26 | 0.20 | 0.59 |
| 13 | T1043-D1 | 148 | 0.16 | 0.18 | 0.18 | 0.22 | 0.27 | 0.22 |
| 14 | T1046s1-D1 | 72 | 0.59 | 0.72 | 0.71 | 0.72 | 0.73 | 0.71 |
| 15 | T1049-D1 | 134 | 0.74 | 0.74 | 0.68 | 0.65 | 0.74 | 0.64 |
| 16 | T1064-D1 | 92 | 0.48 | 0.22 | 0.26 | 0.24 | 0.26 | 0.25 |
| 17 | T1074-D1 | 132 | 0.29 | 0.56 | 0.58 | 0.42 | 0.58 | 0.46 |
| 18 | T1080-D1 | 133 | 0.42 | 0.48 | 0.41 | 0.34 | 0.26 | 0.40 |
| 19 | T1082-D1 | 75 | 0.37 | 0.51 | 0.70 | 0.55 | 0.64 | 0.55 |
| 20 | T1090-D1 | 189 | 0.68 | 0.73 | 0.65 | 0.68 | 0.57 | 0.59 |
